# Development of a tractable model system to mimic wood-ageing of beer on a lab scale

**DOI:** 10.1101/2022.03.11.483928

**Authors:** Sofie Bossaert, Tin Kocijan, Valérie Winne, Filip Van Opstaele, Johanna Schlich, Beatriz Herrera-Malaver, Kevin J. Verstrepen, Gert De Rouck, Bart Lievens, Sam Crauwels

## Abstract

Wood-ageing of conventionally fermented beers is gaining increased attention in the production of sour beers with a noteworthy balance between sourness, wood aroma and flavour complexity. Besides the extraction of wood-derived compounds into the beer, wood-aged sours owe their layered flavour profile to the activity of a variety of ‘wild’ microorganisms that reside in the barrels or that emerge from the brewing or maturation environment. However, until now wood-ageing of craft beers largely remains a process of trial and error that often generates unexpected or undesirable results. Therefore, to better understand the process and develop control strategies to improve the consistency, predictability and overall quality of the resulting beer, more insight is needed into the interactions between the wood, the microorganisms and the maturing beer. Nevertheless, as studying these interactions on an industrial scale is highly challenging, the objective of this study was to develop a reproducible and easy-to-manipulate experimentally tractable system that can be used to study wood-ageing of beer on a lab scale. Barrel-ageing was mimicked in a 0.5 liter glass jar filled with beer and closed off by a wooden disk. Furthermore, the system was equipped with a synthetic community composed of four bacterial species (*Acetobacter malorum, Gluconobacter oxydans, Lactobacillus brevis* and *Pediococcus damnosus*) and four fungal species (*Brettanomyces bruxellensis, Candida friedrichii, Pichia membranifaciens* and *Saccharomyces cerevisiae*) that represented key microbes previously identified in wood-ageing experiments with 225-liter barrels. In order to test the hypothesis that the barrel-ageing process of beer can be replicated in the simplified *in-vitro* system, the system was subjected to 60 days of ageing and microbial community dynamics and beer chemistry were compared with a 38-week industrial barrel-ageing experiment using the same beer. Beer samples were collected at regular time points and subjected to both qPCR assays targeting the eight selected species and chemical analysis. Results revealed that *in vitro* ageing showed similar trends in the temporal dynamics of the microbial populations and beer chemistry as those observed during 38-weeks of barrel-ageing in 225-liter barrels. Furthermore, results were found to be highly reproducible. Altogether, the *in-vitro* system was found to be a robust and reproducible system that has great potential to perform more in-depth research about the intricate interactions between microbes, wood and maturing beer and to develop control strategies to improve the consistency, predictability and overall quality of the resulting beer.

## 1. Introduction

Wood-ageing is an indispensable tool for the production of many alcoholic beverages like whisky, wine, port and traditional beers like lambics and Flanders red ales, because it has positive effects on the flavour and colour of the beverage ageing inside (Cantwell and Bouckaert, 2016; De Roos and De Vuyst, 2019; De Rosso *et al*., 2009). This is also the reason why the technique is gaining increased attention in the production of sour beers obtained from conventionally fermented beer. In this case, following conventional fermentation, beers are aged for a long period of time (often up to several months, one year, or more) in wooden barrels leading to beers with a noteworthy balance between sourness, aroma and flavour complexity (Bossaert *et al*., 2019). Besides the extraction of wood-derived compounds into the beer, wood-aged sours owe their layered flavour profile to the activity of a variety of ‘wild’ microorganisms that reside in the barrels and/or the brewing or maturation environment. However, until now wood-ageing of craft beers largely remains a process of trial and error, which does not always lead to a pleasurable balance between sourness and wood characteristics, but instead it may lead to unexpected, undesirable or even unpalatable results. Therefore, in order to better understand the process and develop process control strategies to improve the consistency, predictability and overall quality of the resulting beer, more insight is needed into the interactions that take place between the environment, the wood, the microorganisms and the maturing beer.

Previous experiments on an industrial scale (Bossaert *et al*., 2021a; 2021b) have revealed temporal patterns in the microbial community structure and beer chemistry throughout the wood-ageing process, and indicated that these temporal dynamics are most probably determined by beer properties like ethanol level and bitterness, wood properties, and environmental characteristics. However, more fundamental in-depth studies under controlled conditions are required to investigate the true effects of these parameters. Nevertheless, challenges arise in the execution of such studies in an accurate and statistically meaningful manner as large quantities of wooden barrels should be purchased, large volumes of beer are needed to fill the barrels, and the barrels should be monitored over a long period of time. Additionally, fluctuating environmental conditions and the variability and complexity of natural microbial communities interfering with the process complicate the experiments. One way to surpass these challenges is by using a lab-scale, experimentally tractable *in-vitro* system based on a number of key microorganisms, that simulates conditions of industrial barrel-ageing. Such small-scale ecosystems can be readily incubated under controlled conditions, and can easily be manipulated to evaluate the impact of diverse parameters of interest. Recently, similar model systems have been developed to assess the mechanisms of microbial community assembly and function *in vitro* (Cosetta *et al*., 2020; Jia *et al*., 2020; Wolfe *et al*., 2014; Wolfe and Dutton, 2015; Wolfe, 2018). For instance, Wolfe *et al*. (2014) used a synthetic *in-vitro* microbial community of 11 key species to mimic cheese rinds (i.e. the microbial fraction that occurs on the surface during cheese ageing), and showed that microbial community assembly is largely determined by abiotic factors like moisture level. In previous research (Bossaert *et al*., 2021a; 2021b) we provided a detailed view on the temporal dynamics of the microbial community composition in the wood-ageing of beer. Beers were found to undergo a shift from a diverse microbial community to a number of key microorganisms, including lactic acid bacteria like *Lactobacillus brevis* and *Pediococcus damnosus*, acetic acid bacteria like *Acetobacter malorum*, and wild yeasts like *Brettanomyces bruxellensis* and *Pichia membranifaciens* (Bossaert *et al*., 2021a; 2021b). Based on these observations, the objective of this study was to develop an easy-to-manipulate, experimentally tractable system to study the mechanisms that underlie microbial successions and chemical changes during barrel-ageing of beer. An ideal model system should be a culture-based system which is simpler than natural communities, yet still exhibits patterns of community formation and dynamics that are representative of those observed *in situ*. Further, the microbial communities involved should assemble in a defined and reproducible way under given experimental conditions. To accomplish this research goal, first a culture collection was established by isolating microbial strains from diverse wood-aged beers, including those from our previous studies (Bossaert *et al*., 2021a; 2021b). Next, based on our previous findings (Bossaert *et al*., 2021a; 2021b) microbial communities were reconstructed by combining key microbes, including four bacterial species (*A. malorum, Gluconobacter oxydans, L. brevis* and *P. damnosus*) and four fungal species (*B. bruxellensis, Candida friedrichii, P. membranifaciens* and *Saccharomyces cerevisiae*). The synthetic community was pitched into experimental microcosm systems mimicking industrial wood-ageing on a lab scale, and incubated for 60 days under controlled conditions. At regular time points, the microbial community dynamics were monitored by species-specific quantitative real-time PCR (qPCR) assays, and the beer chemistry was assessed at the same time points. Obtained results were compared to wood-ageing of the same beer in wooden barrels under industrial conditions (Bossaert *et al*., 2021a).

## 2. Materials and methods

### 2.1 Isolation and community reconstruction

For the isolation of microbes, several samples from diverse wood-aged beers that were monitored throughout a 38-week industrial maturation process, including those from our previous studies (Bossaert *et al*., 2021a; 2021b), were plated on a number of cultivation media (Table 1). These media included Plate Count Ager (PCA, for bacteria in general), de Man-Rogosa-Sharpe Agar (MRSA, for lactic acid bacteria), and Acetic Acid Medium agar (AAM, for acetic acid bacteria), each supplemented with 200 ppm cycloheximide to inhibit fungal growth. Furthermore, samples were plated on Yeast extract Peptone Dextrose agar (YPD, for fungi in general) supplemented with 100 ppm chloramphenicol to inhibit bacterial growth, and Wallerstein Laboratory Nutrient agar (WLN, for cycloheximide-resistant fungi) supplemented with 100 ppm chloramphenicol and 50 ppm cycloheximide to inhibit growth of bacteria and cycloheximide-sensitive fungi, respectively. All plates were incubated aerobically at 30°C for 14 days, except for MRSA which was incubated in anaerobic conditions. Per plate, two distinct colonies (if available) of every morphotype were purified and stored at -80°C in 25% glycerol. Following identification, isolates belonging to all key species previously linked with the barrel-ageing process of conventionally fermented beer were retained for further study (Bossaert *et al*., 2021a; 2021b). These included members of the bacteria *A. malorum, G. oxydans, L. brevis* and *P. damnosus*, as well as members of the yeast species *B. bruxellensis, C. friedrichii, P. membranifaciens* and *S. cerevisiae*. In contrast to *G. oxydans* and *C. friedrichii* which, amongst several others, mainly occur at the beginning of the process, all other species were found to dominate the microbial community over the course of maturation (Bossaert *et al*., 2021a; 2021b; De Roos *et al*., 2018). Identifications were performed by PCR amplification and sequencing of the 16S ribosomal RNA (rRNA) gene (bacteria) and the ITS1-5.8S rRNA-ITS2 region or the D1-D2 region of the large subunit rRNA gene (fungi), followed by a sequence homology search using BLAST against type strains in GenBank as described previously (Jacquemyn *et al*., 2013). For fungi, BLAST searches were also performed against the entire GenBank database.

**Table 1:** Cultivation media used in this study.

| Name medium | Main targets | Ingredients | Concentration | Supplier |
| --- | --- | --- | --- | --- |
| PCA + CY | Aerobic bacteria | PCA<br>Cycloheximide <sup>a</sup> | 17.5 g/l<br>200 mg/l | Oxoid, Thermo Fisher Scientific<br>Sigma-Aldrich |
| MRSA + CY | Lactic acid bacteria | MRSA<br>Cycloheximide <sup>a</sup> | 52 g/l<br>200 mg/l | Oxoid, Thermo Fisher Scientific<br>Sigma-Aldrich |
| AAM + CY <sup>b</sup> | Acetic acid bacteria | Yeast extract<br>Bactopeptone<br>D-glucose <sup>c</sup><br>Agar<br>100% ethanol <sup>d</sup><br>100% acetic acid <sup>d</sup><br>1 N HCl <sup>c</sup><br>Cycloheximide <sup>a</sup> | 8 g/l<br>15 g/l<br>10 g/l<br>20 g/l<br>5 g/l<br>3 g/l<br>28 ml/l (to reach pH 3.5)<br>200 mg/l | Oxoid, Thermo Fisher Scientific<br>BD Biosciences<br>Sigma-Aldrich<br>Oxoid, Thermo Fisher Scientific<br>Fisher Chemical, Thermo Fisher Scientific<br>VWR Chemicals, BDH Prolabo<br>VWR Chemicals, BDH Prolabo<br>Sigma-Aldrich |
| YPD + CH | Fungi | Yeast extract<br>Bactopeptone<br>D-glucose <sup>c</sup><br>Agar<br>Chloramphenicol <sup>a</sup> | 10 g/l<br>20 g/l<br>20 g/l<br>20 g/l<br>100 mg/l | Oxoid, Thermo Fisher Scientific<br>BD Biosciences<br>Sigma-Aldrich<br>Oxoid, Thermo Fisher Scientific<br>Sigma-Aldrich |
| WLN + CH + CY | Cycloheximide resistant fungi | WL nutrient agar<br>Cycloheximide <sup>a</sup><br>Chloramphenicol <sup>a</sup> | 77.2 g/l<br>50 mg/l<br>100 mg/l | Merck<br>Sigma-Aldrich<br>Sigma-Aldrich |

<sup>a</sup> Dissolved in ethanol and added after autoclaving and cooling of the medium.
<sup>b</sup> Based on Lisdiyanti *et al.* (2001).
<sup>c</sup> Dissolved in demineralized water, autoclaved separately from the rest of the medium and mixed after autoclaving.
<sup>d</sup> Filter-sterilized (Rapid-Flow™ bottle top filter, pore size 0.2 µm, Nalgene™, Thermo Fisher Scientific, Waltham, USA) and added to the rest of the medium after autoclaving.

To distinguish different strains belonging to the same species, isolates were subjected to Random Amplified Polymorphic DNA (RAPD) fingerprinting using the RAPD primers OPC20 (5’-ACTTCGCCAC-3’), OPD19 (5’-CTGGGGACTT-3’) and OPK03 (5’-CCAGCTTAGG-3’) (Operon RAPD 10-mer Kits, Operon Technologies, Alameda, USA). As RAPD fingerprinting of *Saccharomyces* spp. did not yield sufficient distinction between different strains, *Saccharomyces* isolates were subjected to interdelta fingerprinting using the primer set delta12 (5’-TCAACAATGGAATCCCAAC-3’) and delta21 (5’-CATCTTAACACCGTATATGA-3’) (Legras and Karst, 2003). All PCR amplifications were performed in a Bio-Rad T100 thermal cycler in a total volume of 20 µl composed of nuclease-free water (Integrated DNA Technologies (IDT), Leuven, Belgium), 0.5 µM primer (one primer for RAPD; two primers for interdelta fingerprinting) (IDT, Leuven, Belgium), 0.15 mM of each dNTP (Invitrogen, Thermo Fisher Scientific, Waltham, USA), 0.4 µl of Titanium *Taq* polymerase (BD Biosciences, Franklin Lakes, USA), 2 µl of Titanium *Taq* buffer (BD Biosciences, Franklin Lakes, USA), and 1 µl (2 ng) of genomic DNA. For the RAPD method, DNA was denatured for 2 minutes at 94°C, followed by 34 cycles that consisted of a one-minute denaturation step at 94°C, a one-minute primer annealing step at 35°C and a two-minute elongation step at 72°C, followed by a final elongation step of 10 minutes at 72°C. For the interdelta method, DNA was denatured for 4 minutes at 95°C, followed by 5 cycles that contained a 30-second denaturation step at 94°C, a 30-second primer annealing step at 42°C and a 90-second elongation step at 72°C. Next, 30 cycles were run that consisted of 30 seconds of denaturation at 95°C, 30 seconds of primer annealing at 45°C and 90 seconds of elongation at 72°C, followed by a final elongation step of 10 minutes at 72°C. Next, the resulting DNA fragments were separated by loading 10 µl of PCR product onto a 1.5% agarose gel stained with ethidium bromide (Invitrogen, Thermo Fisher Scientific, Waltham, USA) in a 1 × Tris/acetate EDTA (TAE) buffer and performing gel electrophoresis at 135 V for two hours. A 1-kb DNA ladder (Smartladder, Eurogentec, Seraing, Belgium) was added as a reference marker. Band patterns were visualized with UV light in an InGenius 3 gel imager (Syngene™, Cambridge, UK), and isolates were considered to be different from others when one or more polymorphisms were observed across the different fingerprints.

For each species, two isolates showing genetic variation were selected and submitted to a pilot screening to monitor the strains’ growth in beer media with different concentrations of ethanol and hop iso-α-acids, and to evaluate their need for supplementary carbon sources. If for certain species isolates represented the same strain, a second strain (if possible, also obtained from beer) from available culture collections was included. Specifically, strains were subjected to the following media: Jupiler (Anheuser-Busch InBev, Leuven, Belgium) (i.e. pilsner beer with 5.2% Alcohol By Volume (ABV) and 13 ppm of iso-α-acids corresponding to a bitterness level of 17 International Bitterness Units (IBU)); Jupiler with increased ethanol levels (8% and 13.5% ABV) (adjusted using 100% ethanol, Fisher Chemical, Thermo Fisher Scientific, Waltham, USA); dry-hopped Jupiler (using Cascade Cryo Hops^®^ pellets, Yakima Chief Hops, Mont-Saint-Guibert, Belgium) (33 IBU, measured before filter sterilization); Jupiler supplemented with iso-α-acids (33, 36, and 45 IBU, measured before filter sterilization) (Brouwland, Beverlo, Belgium); and Jupiler supplemented with 2% of a commercial mixture of sugars that are commonly found in beer (Belgogluc CF-81, HBV-IMTC, Olsene, Belgium). Prior to the screening, selected strains were cultivated on YPD agar plates (Table 1, without chloramphenicol). Subsequently, cells were collected and washed with physiological water (0.85% NaCl), after which the cell density of the cell suspension was adjusted to an Optical Density at 600 nm (OD_600_) equal to 0.1. Next, 200-µl flat-bottom 96-well plates were filled with 140 µl of filter-sterilized beer medium (Rapid-Flow™ bottle top filter, pore size 0.2 μm, Nalgene™, Thermo Fisher Scientific, Waltham, USA) and 10 µl of the cell suspension in each well. For each strain, two biological replicates were included for every beer medium tested. The plates were incubated for 60 hours at 25°C with continuous shaking in a SpectraMax^®^ ABS Plus spectrophotometer equipped with SoftMax^®^ Pro 7.1 software (Molecular Devices, San Jose, USA), and the OD_600_ was measured every 10 minutes. For each species tested, the strain that showed the best performance under the different conditions and that yielded the most consistent results was selected for further experiments.

### 2.2 Development of qPCR assays

For each selected strain, a species-specific PrimeTime™ qPCR probe assay was developed, enabling its accurate detection and quantification. Assays consisted of two primers and a PrimeTime™ double-quenched 5’ 6-FAM/ZEN/3’ IBFQ probe, and were designed from the 16S rRNA gene (bacteria) or the ITS1-5.8S rRNA-ITS2 region (fungi) of the target strains according to the manufacturer’s instructions (IDT, Leuven, Belgium). Briefly, for each strain, a BLAST analysis of the 16S rRNA gene or the ITS1-5.8S rRNA-ITS2 region was performed against type materials in GenBank and the entered sequence was aligned against the most related BLAST results, representing both target and non-target sequences, using the NCBI Multiple Sequence Alignment (MSA) Viewer (version 1.14.0). Alignments were then screened for exploitable differences between the target and non-target sequences, and different species-specific primer sets and probes were designed using the IDT PrimerQuest™ tool (Owczarzy *et al*., 2008), as described by the supplier (IDT, Leuven, Belgium). Primers and probes were designed to have a melting temperature between 60 and 64°C and a GC content between 45 and 60%, with a maximum amplicon length of 120 bp. Further, primers were developed in such a way that mismatches with non-target sequences were located at the 3’ end of the primers and that formation of hairpin structures and primer dimers is avoided (assessed by the Oligonucleotide Properties Calculator ‘OligoCalc’ (Kibbe, 2007)). Next, specificity of the primers and probes was verified by a BLAST analysis against type strains in GenBank, and the most promising primer-probe sets were retained for further tests in the laboratory. Specificity of the assays was assessed by subjecting each assay to 1 ng of genomic DNA from every strain selected. Additionally, for each assay, the limit of quantification (LOQ, defined as the lowest amount of target DNA in the linear range of the assay that can be determined quantitatively) was determined using a 10-fold dilution series of pure genomic DNA (ranging from 5 ng/µl to 5 × 10^−7^ ng/µl) and a 10-fold dilution series of the target DNA fragment (created by PCR amplification) (ranging from 5 ng/µl to 5 × 10^−10^ ng/µl). Furthermore, the linear dynamic range was determined. qPCR reactions were carried out in a StepOnePlus Real-Time PCR system (Applied Biosystems, Thermo Fisher Scientific, Waltham, USA) according to the PrimeTime^®^ Gene Expression Master Mix (IDT, Leuven, Belgium) protocol using an annealing temperature of 62°C for all assays. For each sample, the threshold cycle (C_t_) was calculated with the supplied StepOne™ software, and the baseline was set automatically above any noise. All qPCRs were performed in duplicate, and in every run a negative control was included in which the template DNA was replaced by sterile water. In all analyses, samples were considered positive when the measured C_t_ value was lower than that of the blank.

### 2.3 Model system

Conditions of industrial-scale barrel-ageing of beer were mimicked by the model system shown in Fig. 1. Briefly, the system was based on a 0.5-liter weck jar (diameter: 9.5 cm, height: 8.5 cm; Ikea, Zaventem, Belgium) of which the lid was replaced by a wooden disk with a thickness of 3.5 cm. The wood was supplied by Garbellotto Spa (Pordenone, Italy) and represented new (unused) oak wood of NIR category ‘sweet’ (Bossaert *et al*., 2021a; 2021b) that would otherwise be used for barrel manufacturing. To simulate beer ageing in industrial-scale barrels, the ratio between the beer volume and the wood contact surface area was set to the ratio found in 225-liter barrels. Further, the pressure inside the jars was released via a water lock (diameter of the opening at the bottom: 9 mm; Brouwland, Beverlo, Belgium) that was put through a 13-mm hole in the bottom of the jar with a silicon plug (Fig. 1). Before use, the glass jars, sealing rings, metal clasps, silicon plugs and water locks were thoroughly cleaned and disinfected using 70% ethanol. Next, the jars were filled with a filter-sterilized (Rapid-Flow™ bottle top filter, pore size 0.2 μm, Nalgene™, Thermo Fisher Scientific, Waltham, USA) 0.1% citric acid solution (Vinoferm, Brouwland, Beverlo, Belgium) and incubated upside down for five days at 20°C to saturize the wood pores and to remove the most pungent tannins. The same pre-treatment is recommended when using new oak barrels in industrial barrel-ageing, and has also been performed in our earlier experiments (Bossaert *et al*., 2021a; 2021b). Finally, the jars were emptied, again disinfected with 70% ethanol, and filled with 500 ml beer. Similar beer was used as in Bossaert *et al*. (2021a), i.e. a top-fermented blond beer with an alcohol content of 8.88% ABV and an intermediate bitterness corresponding to an iso-α-acid concentration of 19.85 ppm. The beer was produced by brewing wort to an initial density of 18.8°P using pilsner malt, supplemented with high maltose syrup (Bossaert *et al*., 2021a). Further, Belgian Target hop was used, no dry-hopping was performed, and primary fermentation was executed by *S. cerevisiae* for five days at 20°C and was not cold crashed (Bossaert *et al*., 2021a). The beer was filter-sterilized (Rapid-Flow™ bottle top filter, pore size 0.2 μm, Nalgene™, Thermo Fisher Scientific, Waltham, USA) before use in the experiment *in vitro*.

**Figure 1:**
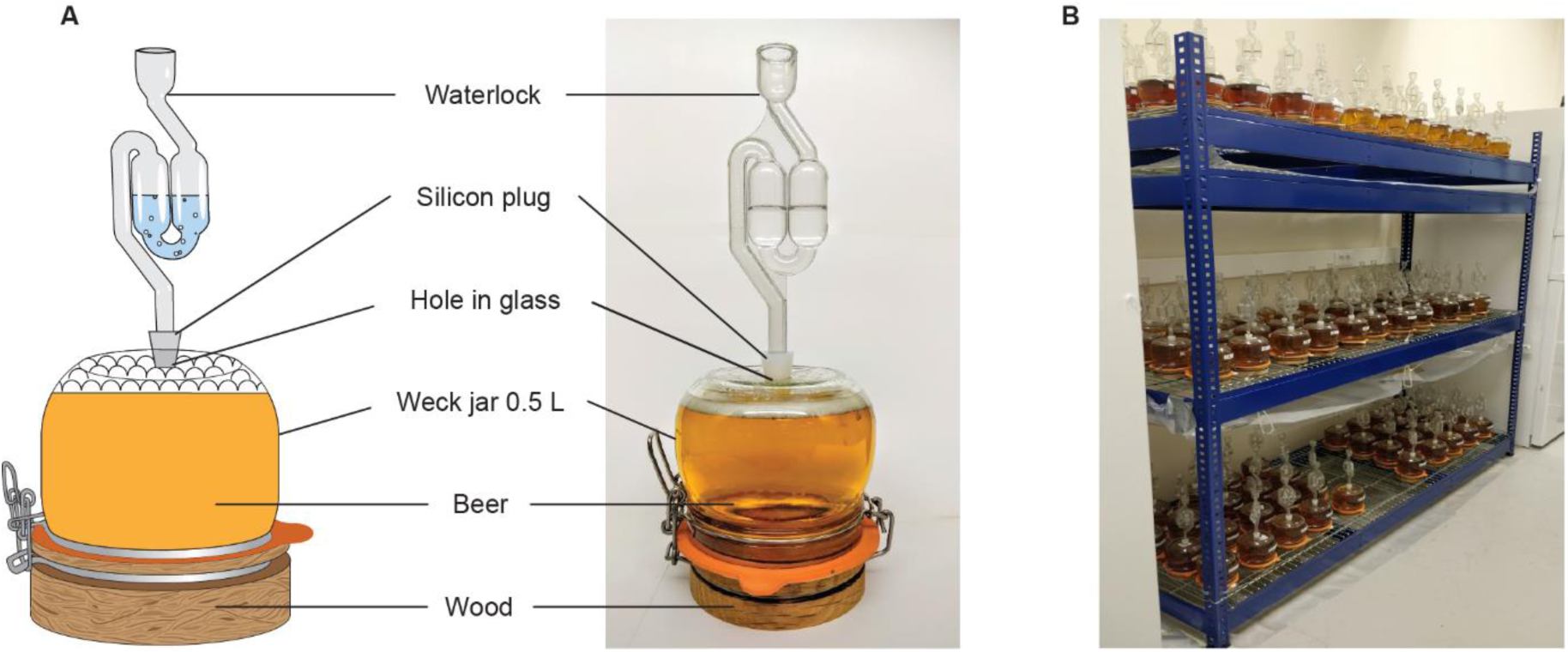
(A) Illustration of the *in-vitro* system used in this study and (B) the incubation of the jars on a rack with a grid to allow oxygen ingress. The design is based on a 0.5-liter weck jar in which the lid is replaced by a wooden disk made from new (European) oak that would otherwise be used for barrel manufacturing. To mimic the extraction of wood-derived compounds and the oxygen ingress rate found in industrial-scale barrels, the ratio between the beer volume and the wood surface area is the same as in 225-liter barrels. The pressure inside the jars is released via a water lock that is positioned through a hole in the glass jar.

To test whether the patterns of microbial community composition and succession observed in our previous experiments (Bossaert *et al*., 2021a; 2021b) can be recapitulated in a simple *in-vitro* system, jars were pitched with a synthetic microbial community composed of the eight selected strains. Prior to composing the synthetic microbial community, individual strains were cultivated at 30°C on YPDF agar that contained 1% yeast extract (Oxoid, Thermo Fisher Scientific, Waltham, USA), 2% bactopeptone (BD Biosciences, Franklin Lakes, USA), 1% D-glucose (Sigma-Aldrich, St. Louis, USA), 1% fructose (Acros Organics, Thermo Fisher Scientific, Waltham, USA) and 2% agar (Agar bacteriological no. 1, Oxoid, Thermo Fisher Scientific, Waltham, USA). *Lactobacillus brevis* and *P. damnosus* were cultivated in anaerobic conditions, while all other strains were cultivated aerobically. Following incubation, for each strain cells were collected, washed, and resuspended in sterile physiological water (0.85% NaCl) before combining them and pitching the resulting community to the beer to reach a final cell density of 10 colony forming units (CFU)/ml per strain in the beer, using a total pitching volume of 1 ml. Jars in which no microbes were pitched, but which also contained the wood were included as a negative control. All jars were incubated statically in the dark for 60 days in a conditioned room with an ambient temperature of 19.69°C ± 0.01°C and a relative humidity equal to 52.07% ± 0.04% (Table S1, Supplementary Information).

### 2.4 Sample collection

Samples were taken immediately after filling and pitching the jars (i.e. at day 0), and at day 10, 20, 30 and 60, by sacrificing three jars per time point (i.e. three biological replicates). Hence, this resulted in a total of 15 samples. The negative controls were analysed at the end of the experiment, and also represented three jars. Sampling was performed by thoroughly shaking the beer, followed by removing the water lock and draining the entire homogenized beer volume through the hole in the bottom of the jar. The collected beer was then centrifuged at 3,500 × g for 15 minutes at 4°C, and obtained cell pellets and supernatants were preserved at -20°C for microbiological and chemical analyses, respectively (further referred to as samples from the ‘*in-vitro* experiment’). As a reference, previously studied samples taken from ‘beer 3’, matured in 225-liter oak barrels of the NIR-category ‘sweet’ were again included in this study (Bossaert *et al*., 2021a). In this case, samples were taken after 0, 2, 12, and 38 weeks of maturation and stored at -20°C as described previously (Bossaert *et al*., 2021a) (further referred to as samples from the ‘*in-situ* experiment’).

### 2.5 Microbiological analyses

For each sample, genomic DNA was extracted in duplicate from 500 µl of the cell pellet that was dissolved in 5 ml of physiological water (0.85% NaCl) according to the DNA extraction procedure described in Lievens *et al*. (2003). Following mixing of both DNA replicates, qPCR amplification was performed in duplicate using each of the PrimeTime™ qPCR probe assays developed, according to the procedures outlined above. In each run, two negative controls were included in which sterile nuclease-free water was used instead of template DNA. For each assay, bacterial 16S rRNA gene or fungal ITS copy numbers were calculated via standard curves obtained from a 10-fold amplicon dilution series that was included in each run. The qPCR quantification cut-off was based on the C_t_-values obtained for the negative controls and was set to a C_t_-value of 34 for all assays. In subsequent data analysis, gene copy numbers were set to zero when the C_t_ value was equal to or higher than 34.

### 2.6 Chemical analyses

Chemical analyses were performed according to protocols described in Bossaert *et al*. (2021a). Briefly, for each sample, wood-derived aroma compounds were quantified via Head Space – Solid Phase Micro Extraction – Gas Chromatography – Mass Spectrometry (HS-SPME-GC-MS), while Head Space – Gas Chromatography – Flame Ionization Detector (HS-GC-FID) was used to measure the concentration of fermentation products (esters and higher alcohols). Additionally, the pH and concentrations of carbohydrates, organic acids and total polyphenols were assessed using a Gallery Plus Beermaster (Thermo Fisher Scientific, Waltham, USA), and the ethanol content was measured with an Alcolyzer beer ME (Anton Paar GmbH, Graz, Austria). A detailed overview of the protocols used for sample preparation and chemical analyses can be found in Table S2 (Supplementary information).

### 2.7 Data visualization and statistical analyses

Permutational multivariate analysis of variance (perMANOVA) was performed to test whether there was a significant difference in gene copy numbers per µl DNA for the eight strains studied, between the maturation experiment *in vitro* and *in situ*. The analysis was performed using data from the start of the experiment (day 0) and the end of the experiment (day 60 for the *in-vitro* experiment, and week 38 for the *in-situ* experiment). Additionally, within each experiment, perMANOVA was used to test whether maturation time had a significant effect on the gene copy numbers measured for the different strains. All perMANOVAs were calculated using the adonis function of the ‘vegan’ R-package (v3.6.1) (Oksanen *et al*., 2019). Bray-Curtis distances of the gene copy numbers per µl DNA were used to create a non-metric multidimensional scaling (NMDS) plot that visualizes the differences in the bacterial and fungal community structure throughout the wood maturation process in both the *in-vitro* and *in-situ* experiment. Therefore, the MDS function of the ‘vegan’ package in R was used (Oksanen *et al*., 2019). Furthermore, line plots were constructed, for each strain, showing the temporal dynamics in gene copy numbers per µl DNA averaged over the different biological replicates per time point (*n* = 3 for the experiment *in vitro*, and *n* = 2 for the experiment *in situ*), and the associated data variation is shown as the standard error of the mean. Likewise, perMANOVA was used to assess significant differences in beer chemistry at the start and at the end of the maturation period, as well as to assess the impact of the maturation time within each experiment. Further, a Principal Component Analysis (PCA) was carried out with the scaled chemical parameters, using the ‘stats’ package in R (R Core Team, 2019), to reveal differences in beer chemistry between samples of both experiments, and samples taken at different time points during the maturation process. Additionally, heatmaps were created with MS Excel, showing the temporal changes in chemical parameters as the ratio between the concentration measured at each time point and the concentration at the start (day 0). Likewise, the ratio between the concentrations measured at the last time points in both experiments was calculated to reveal differences in end concentration for each parameter. Welch t-tests were performed (using the ‘stats’ package in R; R Core Team, 2019) to reveal whether the end concentrations of each parameter were significantly different from one another. Finally, for the experiment *in vitro*, the ratio between the concentrations of chemical parameters in the negative controls at day 60 and their concentration at day 0 was calculated, as well as the ratio between the concentrations measured in samples inoculated with the synthetic microbial community at day 60 and in the negative controls at day 60, to visualize the effect of the maturation in contact with wood, and the effect of the microbial community on each chemical parameter, respectively. Moreover, Welch t-tests were performed to assess whether the concentrations of chemical compounds in the numerator and the denominator of the visualized ratios were significantly different.

## 3. Results

### 3.1 Isolation and characterization of key microbes

In total, 751 microbial isolates were obtained from the beer samples plated, and 58 isolates were identified as *A. malorum* (7 isolates), *G. oxydans* (9), *L. brevis* (3), *P. damnosus* (5), *B. bruxellensis* (17), *C. friedrichii* (8), *P. membranifaciens* (1), or *S. cerevisiae* (8). Among these, based on genetic fingerprinting, two different strains per species were selected to evaluate their growth in different beer media. For *G. oxydans, C. friedrichii* and *P. membranifaciens*, however, only one strain was obtained. Therefore, for these species, an additional strain obtained from available culture collections was included, i.e. *G. oxydans* LMG 1519 (LMG, Ghent, Belgium), *C. friedrichii* MUCL 27715 (MUCL, Ottignies-Louvain-la-Neuve, Belgium) and *P. membranifaciens* ST01.13/005 (PME&BIM culture collection). For each species, the best performing strain showing the most consistent results, with a high tolerance to ethanol and hop compounds, and which grew well without the need of supplementary carbon sources, was selected for further experiments (Table 2).

**Table 2:**
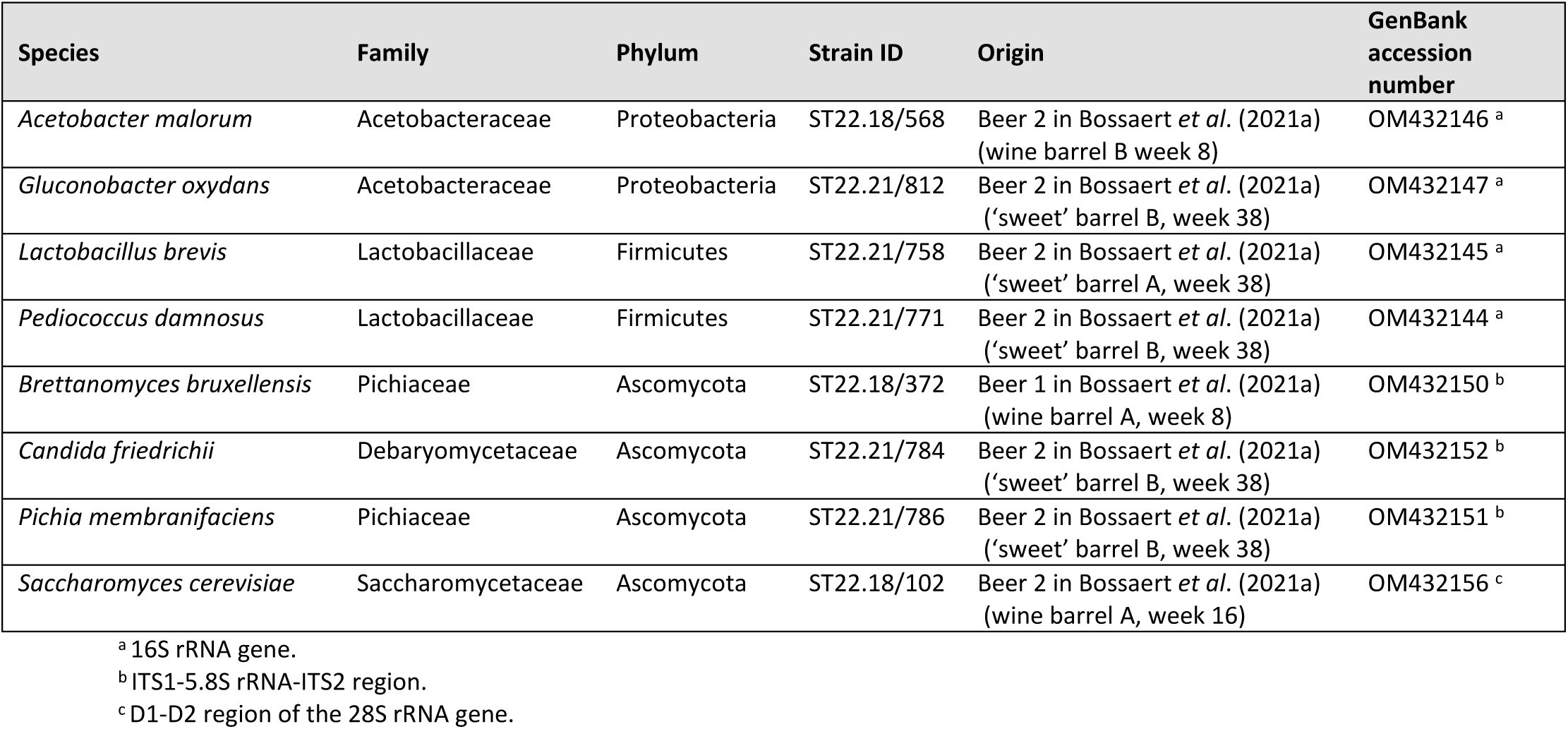
Strains selected for this study.

### 3.2 Development of qPCR assays

For each target strain, multiple primer-probe sets were designed and evaluated *in silico*, among which the most promising combination was retained for further use (Table 3).

**Table 3:**
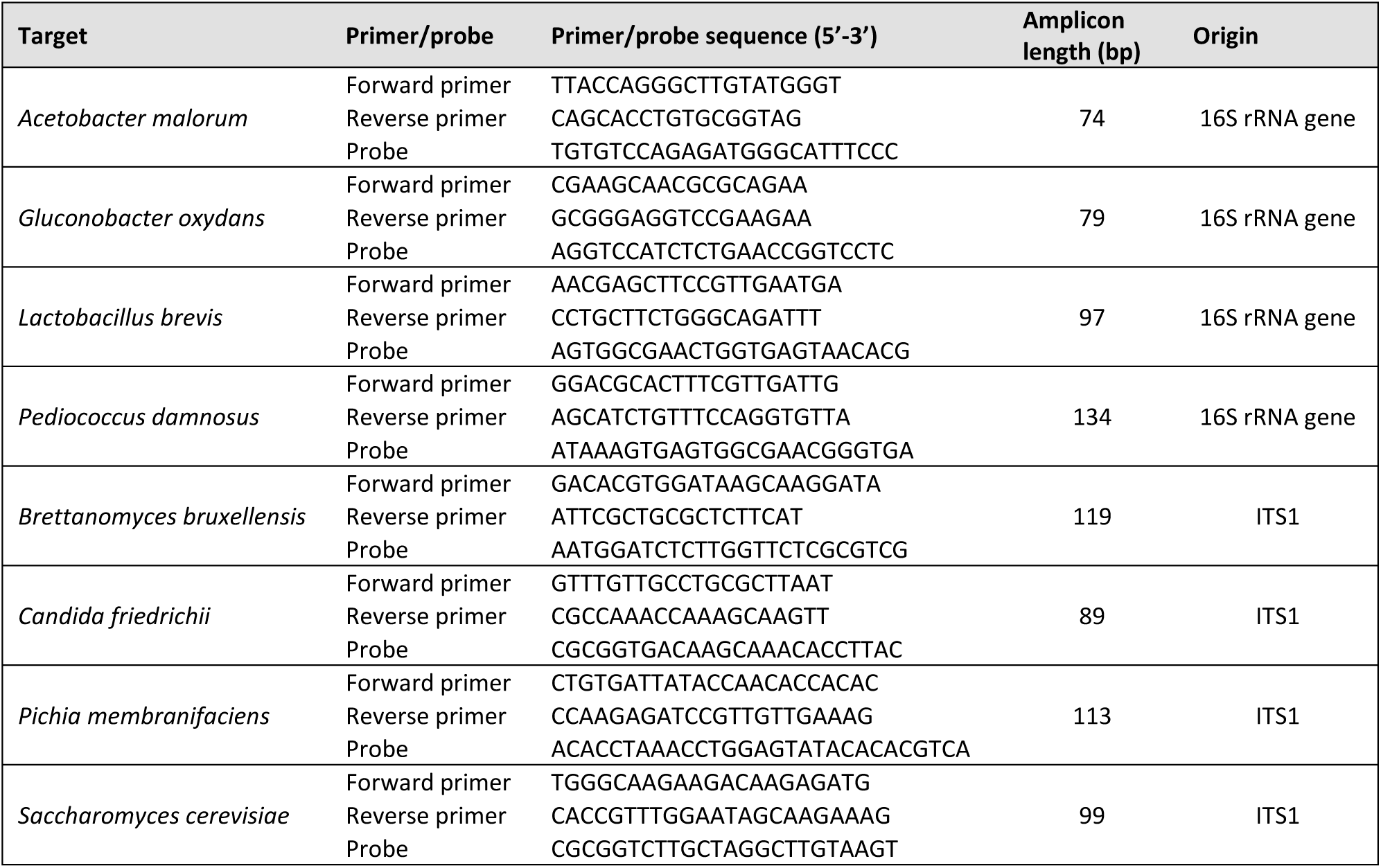
Primers and probes used in this study.

Assessment of the specificity of the different assays using 1 ng of genomic DNA revealed that only the target strains showed a positive response, and that no cross-reaction with any other selected strain was obtained (Table 4).

**Table 4:** Specificity assessment of each qPCR assay against all strains selected for this study ^a^.

| Species | Strain ID | Assay target species |  |  |  |  |  |  |  |
| --- | --- | --- | --- | --- | --- | --- | --- | --- | --- |
|  |  | *A. malorum* | *G. oxydans* | *L. brevis* | *P. damnosus* | *B. bruxellensis* | *C. friedrichii* | *P. membranifaciens* | *S. cerevisiae* |
| *Acetobacter malorum* | ST22.18/568 | + | - | - | - | - | - | - | - |
| *Gluconobacter oxydans* | ST22.21/812 | - | + | - | - | - | - | - | - |
| *Lactobacillus brevis* | ST22.21/758 | - | - | + | - | - | - | - | - |
| *Pediococcus damnosus* | ST22.21/771 | - | - | - | + | - | - | - | - |
| *Brettanomyces bruxellensis* | ST22.18/372 | - | - | - | - | + | - | - | - |
| *Candida friedrichii* | ST22.21/784 | - | - | - | - | - | + | - | - |
| *Pichia membranifaciens* | ST22.21/786 | - | - | - | - | - | - | + | - |
| *Saccharomyces cerevisiae* | ST22.18/102 | - | - | - | - | - | - | - | + |

<sup>a</sup> + : positive detection, - : negative detection. Results were considered positive when obtained $C_t$ values were lower than that of the blanks ( $C_t = 34$ ).

When testing a 10-fold dilution series of genomic DNA, the qPCRs were found to have a linear dynamic range of at least four orders of magnitude and LOQ values ranging from 2.6 to 75.2 fg genomic DNA per µl. When a 10-fold dilution series of PCR amplicons was subjected to the assays, a linear dynamic range of at least nine orders of magnitude was obtained, with LOQ values ranging between 0.81 and 1.88 log gene copies per µl (Table 5; Fig. S1 and S2, Supplementary Information).

**Table 5:** Quantification characteristics of the qPCR assays developed in this study.

| Target | Genomic DNA (DNA concentration, fg/$\mu\text{l}$) | | Amplicons (log gene copy numbers per $\mu\text{l}$ DNA) | |
| --- | --- | --- | --- | --- |
|  | LOQ <sup>a</sup> | Dynamic range <sup>b</sup> | LOQ <sup>a</sup> | Dynamic range <sup>b</sup> |
| *Acetobacter malorum* | 25.0 | 25.0 – 5.9 $\times 10^5$ | 1.31 | 1.31 – 9.83* |
| *Gluconobacter oxydans* | 27.5 | 27.5 – 5.0 $\times 10^5$ | 1.32 | 1.32 – 9.74* |
| *Lactobacillus brevis* | 75.2 | 75.2 – 5.4 $\times 10^5$ | 1.46 | 1.46 – 10.71 |
| *Pediococcus damnosus* | 38.9 | 38.9 – 4.7 $\times 10^5$ | 1.62 | 1.62 – 10.59 |
| *Brettanomyces bruxellensis* | 6.9 | 6.9 – 5.8 $\times 10^5$ | 0.81 | 0.81 – 9.63* |
| *Candida friedrichii* | 21.8 | 21.8 – 5.5 $\times 10^5$ | 1.29 | 1.29 – 9.73* |
| *Pichia membranifaciens* | 2.6 | 2.6 – 5.6 $\times 10^5$ | 1.24 | 1.24 – 9.67 |
| *Saccharomyces cerevisiae* | 37.2 | 37.2 – 5.8 $\times 10^5$ | 1.88 | 1.88 – 10.71 |

<sup>a</sup> Limit of quantification (LOQ), defined as the lowest quantity of DNA (DNA concentration or gene copy number) in the linear range of the assay that can be measured with a cut-off value of $C_t = 34$ (corresponding to the lowest $C_t$ value obtained for the blanks).
<sup>b</sup> Dynamic range, defined as the range of DNA concentrations or the range of gene copy numbers in which linearity is maintained. The lower end of the range is equal to the LOQ, and the higher end of the range is equal to the highest tested DNA concentration or number of gene copies. However, when the highest tested DNA quantity did not fall within the linear range of the assay, then the second highest tested DNA quantity is reported (indicated with an asterisk).

### 3.3 Reconstruction of microbial community dynamics

To assess whether the patterns of microbial community composition observed *in situ* can be reiterated in the *in-vitro* system, population dynamics of the eight species selected were monitored by qPCR throughout the maturation process of 60 days for the *in-vitro* experiment and 38 weeks for the *in-situ* experiment (Table S3, Supplementary Information). PerMANOVA revealed that gene copy numbers of the different bacteria and fungi were not significantly different between both experiments at day 0, nor at the last sampling point (Table 6). PerMANOVA also revealed a significant effect of maturation time on both the bacterial and fungal populations in the *in-vitro* system, but not for the *in-situ* experiment (possibly due to the limited number of replicates in this experiment) (Table 6). NMDS ordination, however, suggested that the bacterial populations (stress = 0.081) changed throughout the course of maturation in both experiments. Furthermore, the NMDS plot indicates that the bacterial populations in both experiments converged to a similar community composition as can be seen from the NMDS plot in which samples taken at the final sampling point (day 60 for the experiment *in vitro*, and week 38 for the experiment *in situ*) are plotted closely together (Fig. 2A). For fungi (stress = 0.031), patterns were less similar between both experiments. Specifically, fungal populations in the barrels remained relatively stable over time, while the samples taken at different time points in the *in-vitro* system are clearly separated (Fig. 2B). Importantly, biological replicates were all plotted closely together in the NMDS plot, reinforcing the robustness and reproducibility of the *in-vitro* system under study (Fig. 2).

**Table 6:** Results of permutational multivariate analysis of variance (perMANOVA) ^a^ comparing the abundance of the set of eight bacterial ^b^ and fungal ^c^ species between the experiment *in vitro* and the experiment *in situ*, as well as the effect of maturation time on the bacterial and fungal populations within each experiment.

| Copy numbers at the starting point (23 permutations) <sup>d</sup> |  |  |  |  |  |  |
| --- | --- | --- | --- | --- | --- | --- |
| Bacteria |  |  |  | Fungi |  |  |
| Factors | df | F | p | df | F | p |
| Treatment | 1 | 15.29 | 0.25 n.s. | 1 | 30.836 | 0.25 n.s. |
| Copy numbers at the end point (119 permutations) <sup>e</sup> |  |  |  |  |  |  |
| Bacteria |  |  |  | Fungi |  |  |
| Factors | df | F | p | df | F | p |
| Treatment | 1 | 8.432 | 0.1 n.s. | 1 | 18.529 | 0.1 n.s. |
| Copy numbers experiment *in vitro* (1,000 permutations) <sup>f</sup> |  |  |  |  |  |  |
| Bacteria |  |  |  | Fungi |  |  |
| Factors | df | F | p | df | F | p |
| Time | 4 | 36.274 | < 0.001 *** | 4 | 24.833 | < 0.001 *** |
| Copy numbers experiment *in situ* (5,039 permutations) <sup>g</sup> |  |  |  |  |  |  |
| Bacteria |  |  |  | Fungi |  |  |
| Factors | df | F | p | df | F | p |
| Time | 3 | 2.598 | 0.1009 n.s. | 3 | 6.0653 | 0.08891 n.s. |

<sup>a</sup> df: degrees of freedom, F: F-statistic, p: p-value (values are statistically significant when $p < 0.05$ ), n.s.: not significant ( $p \geq 0.05$ ), \*\*\* $p < 0.001$ .
<sup>b</sup> Bacterial communities were compared using 16S rRNA gene copy numbers for the four bacteria pitched in the *in-vitro* system: *Acetobacter malorum*, *Gluconobacter oxydans*, *Lactobacillus brevis*, and *Pediococcus damnosus*.
<sup>c</sup> Fungal communities were compared using ITS copy numbers for the four fungi pitched in the *in-vitro* system: *Brettanomyces bruxellensis*, *Candida friedrichii*, *Pichia membranifaciens*, and *Saccharomyces cerevisiae*.
<sup>d</sup> Analysed samples include three biological replicates of the *in-vitro* system at day 0 and one sample of the beer in wooden barrels at day 0, limiting the number of permutations to 23.
<sup>e</sup> Analysed samples include three biological replicates of the *in-vitro* system at day 60 and two samples of the beer in wooden barrels at week 38, limiting the number of permutations to 119.
<sup>f</sup> Analysed samples include three biological replicates for each of the selected time points: day 0, day 10, day 20, day 30, and day 60. The number of permutations was set to 1,000.
<sup>g</sup> Analysed samples include one sample of the beer at week 0, and two biological replicates at each of the other selected time points: week 2, week 12, and week 38. In total, 5,039 permutations were performed.

**Figure 2:**
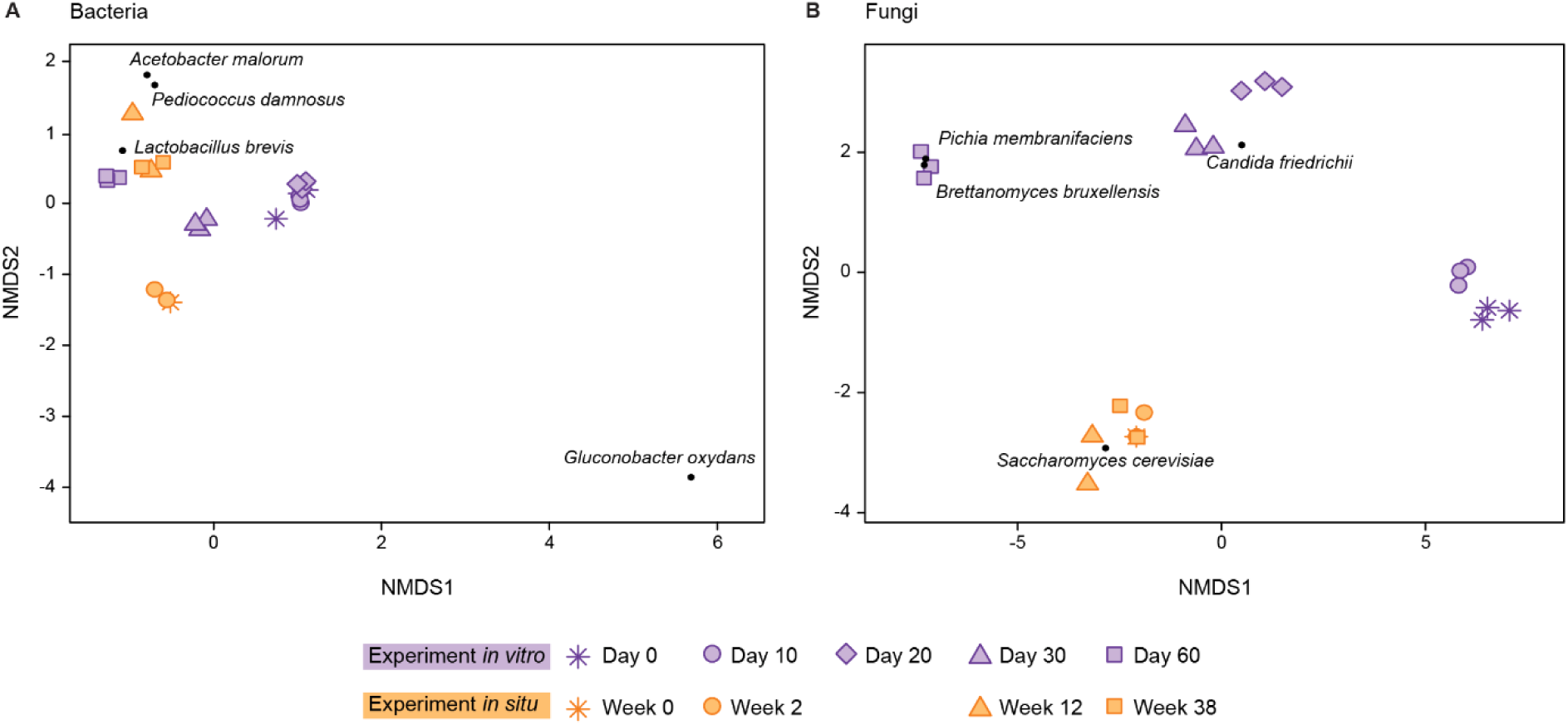
Non-metric multidimensional scaling (NMDS) plot based on Bray-Curtis distances comparing (A) bacterial and (B) fungal populations in both the *in-vitro* and *in-situ* experiment. Plots are based on the number of gene copies per µl DNA obtained for the bacteria *Acetobacter malorum, Gluconobacter oxydans, Lactobacillus brevis* and *Pediococcus damnosus* (stress = 0.081), and the fungi *Brettanomyces bruxellensis, Candida friedrichii, Pichia membranifaciens* and *Saccharomyces cerevisiae* (stress = 0.031). In the *in-vitro* experiment (*n* = 3), microbial dynamics were monitored over 60 days. In the *in-situ* experiment (*n* = 2), microbial dynamics were monitored throughout a maturation period of 38 weeks in 225-liter oak barrels. The same beer was used in both experiments. However, whereas the barrels were filled with non-filtered beer that still contained the *S. cerevisiae* strain from primary fermentation, the beer used in the *in-vitro* experiment was filter-sterilized and inoculated with a synthetic microbial community composed of the eight species mentioned above.

When zooming in on the dynamics of the individual species (Fig. 3), it becomes clear that the 16S rRNA gene copy numbers of *L. brevis* increased considerably over time in both experiments, i.e. from 0.59 ± 0.48 log gene copies per µl DNA at the start of the experiment to 7.29 ± 0.06 log 16S rRNA gene copies per µl DNA at the end of the experiment in the inoculated jars, and from 1.77 to 6.76 ± 0.14 log gene copies per µl DNA in the 225-liter barrels. Likewise, in both experiments, gene copy numbers of *G. oxydans* remained very low. More specifically, in the weck jars gene copy numbers of *G. oxydans* decreased from 2.12 ± 0.04 log gene copies per µl DNA to copy numbers smaller than the detection limit (C_t_ value equal to or higher than 34) in 60 days, while gene copy numbers remained under the detection limit throughout the entire 38-week maturation period in barrels. In contrast, the temporal dynamics in 16S rRNA gene copy numbers of *A. malorum* and *P. damnosus* differed between both experiments. Whereas gene copy numbers of *A. malorum* and *P. damnosus* remained low in the weck jars, both bacterial species were found at higher densities in the barrels, i.e. gene copy numbers changed from below the detection limit for both species to 5.74 ± 0.57 log gene copy numbers of *A. malorum* per µl DNA and to 6.53 ± 0.06 log gene copy numbers of *P. damnosus* per µl DNA at the end of the experiment. Also fungal ITS copy numbers changed throughout maturation. In fact, in the *in-vitro* system, log ITS copy numbers of *B. bruxellensis, P. membranifaciens*, and *S. cerevisiae* increased from below the detection limit for *B. bruxellensis* and *P. membranifaciens* to 7.74 ± 0.02 and 6.85 ± 0.17 log ITS copies per µl DNA after 60 days of maturation, respectively, and from 1.73 ± 0.06 to 5.32 ± 0.08 log ITS copies per µl DNA for *S. cerevisiae*, while ITS copy numbers of *C. friedrichii* were below the detection limit at the start and after 60 days of ageing *in vitro*. Similarly, in the barrels, the ITS copy numbers of *B. bruxellensis, P. membranifaciens* and *S. cerevisiae* increased over the course of 38 weeks of barrel-ageing, although changes in ITS copy numbers were smaller than found in the *in-vitro* experiment. Additionally, similar to what was observed in the *in-vitro* system, ITS copies of *C. friedrichii* remained below the detection limit in the barrels (Fig. 3).

**Figure 3:**
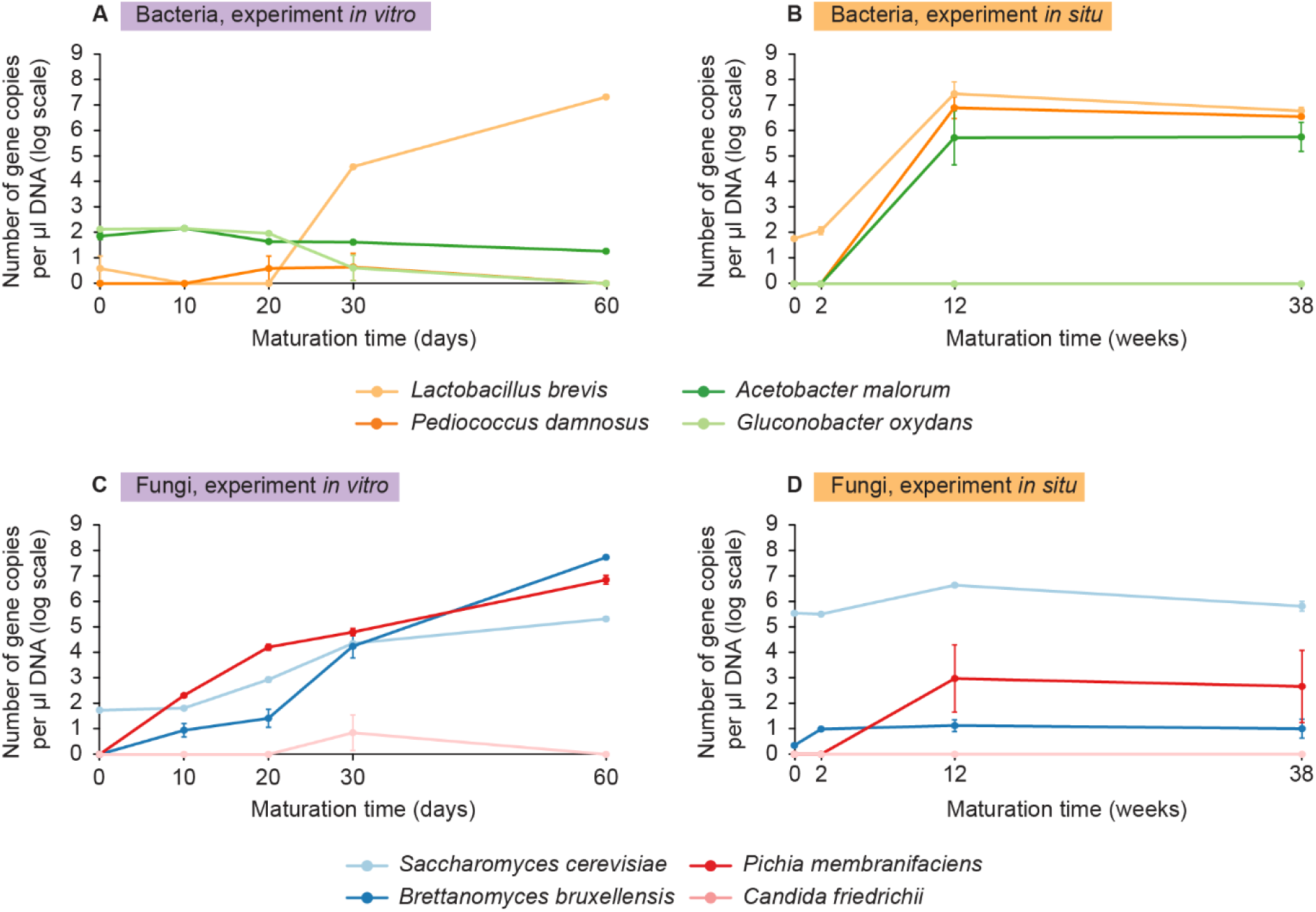
Temporal dynamics in bacterial and fungal community composition for (A and C) the experiment *in vitro* and (B and D) the experiment *in situ*. Plots are based on the number of gene copies per µl DNA obtained for the bacteria *Acetobacter malorum, Gluconobacter oxydans, Lactobacillus brevis* and *Pediococcus damnosus*, and the fungi *Brettanomyces bruxellensis, Candida friedrichii, Pichia membranifaciens* and *Saccharomyces cerevisiae*. In the *in-vitro* experiment (*n* = 3), microbial dynamics were monitored over 60 days. In the *in-situ* experiment (*n* = 2), microbial dynamics were monitored throughout a maturation period of 38 weeks in 225-liter oak barrels. The same beer was used in both experiments. However, whereas the barrels were filled with non-filtered beer that still contained the *S. cerevisiae* strain from primary fermentation, the beer used in the *in-vitro* experiment was filter-sterilized and inoculated with a synthetic microbial community composed of the eight species mentioned above. Gene copy numbers were set to zero when the C_t_ value was equal to or higher than that of the blanks (C_t_ = 34). Error bars represent standard error of the mean.

### 3.4 Reconstruction of dynamics in beer chemistry

PerMANOVA indicated that beer samples from both experiments did not have a significantly different chemical composition at the starting point, nor at the last sampling point. Further, PerMANOVA revealed that maturation time significantly affected the beer chemistry within each experiment (Table 7). The effect of maturation time can also be seen from the PCA ordination plot (Fig. 4), in which samples of different time points are clearly separated by the first PCA axis. Samples taken after 60 days of ageing in the *in-vitro* system and after 38 weeks of ageing in 225-liter barrels were characterized by low levels of sugars and higher levels of wood-related compounds like vanillin, total polyphenols, cis- and trans-3-methyl-4-octanolide, as well as organic acids including lactic acid, acetic acid and propionic acid, and the microbial metabolites 4-ethyl phenol and 4-ethyl guaiacol (Fig. 4).

**Table 7:** Permutational multivariate analysis of variance (perMANOVA) ^a^ comparing beer chemistry between the experiment *in vitro* and the experiment *in situ*, as well as the effect of maturation time on the chemical composition of the beer within each experiment.

| Beer chemistry starting point (23 permutations) <sup>b</sup> |  |  |  |  |
| --- | --- | --- | --- | --- |
| Factors | df | F | p |  |
| Treatment | 1 | 2.9493 | 0.25 | n.s. |
| Beer chemistry end point (119 permutations) <sup>c</sup> |  |  |  |  |
| Factors | df | F | p |  |
| Treatment | 1 | 33.801 | 0.1 | n.s. |
| Beer chemistry experiment *in vitro* (1,000 permutations) <sup>d</sup> |  |  |  |  |
| Factors | df | F | p |  |
| Time | 4 | 3.876 | 0.008991 | ** |
| Beer chemistry barrel experiment *in situ* (5,039 permutations) <sup>e</sup> |  |  |  |  |
| Factors | df | F | p |  |
| Time | 3 | 23.571 | 0.00999 | ** |

<sup>a</sup> df: degrees of freedom, F: F-statistic, p: p-value (values are statistically significant when $p < 0.05$ ), n.s.: not significant ( $p \geq 0.05$ ), \*\* $p < 0.01$ .
<sup>b</sup> Analysed samples include three biological replicates of the *in-vitro* system at day 0 and one sample of the beer in wooden barrels at day 0, limiting the number of permutations to 23.
<sup>c</sup> Analysed samples include three biological replicates of the *in-vitro* system at day 60 and two samples of the beer in wooden barrels at week 38, limiting the number of permutations to 119.
<sup>d</sup> Analysed samples include three biological replicates for each of the analysed time points: day 0, day 10, day 20, day 30, and day 60. The number of permutations was set to 1,000.
<sup>e</sup> Analysed samples include one sample of the beer at week 0, and two biological replicates at each of the other analysed time points: week 2, week 12, week 38. In total, 5,039 permutations were performed.

**Figure 4:**
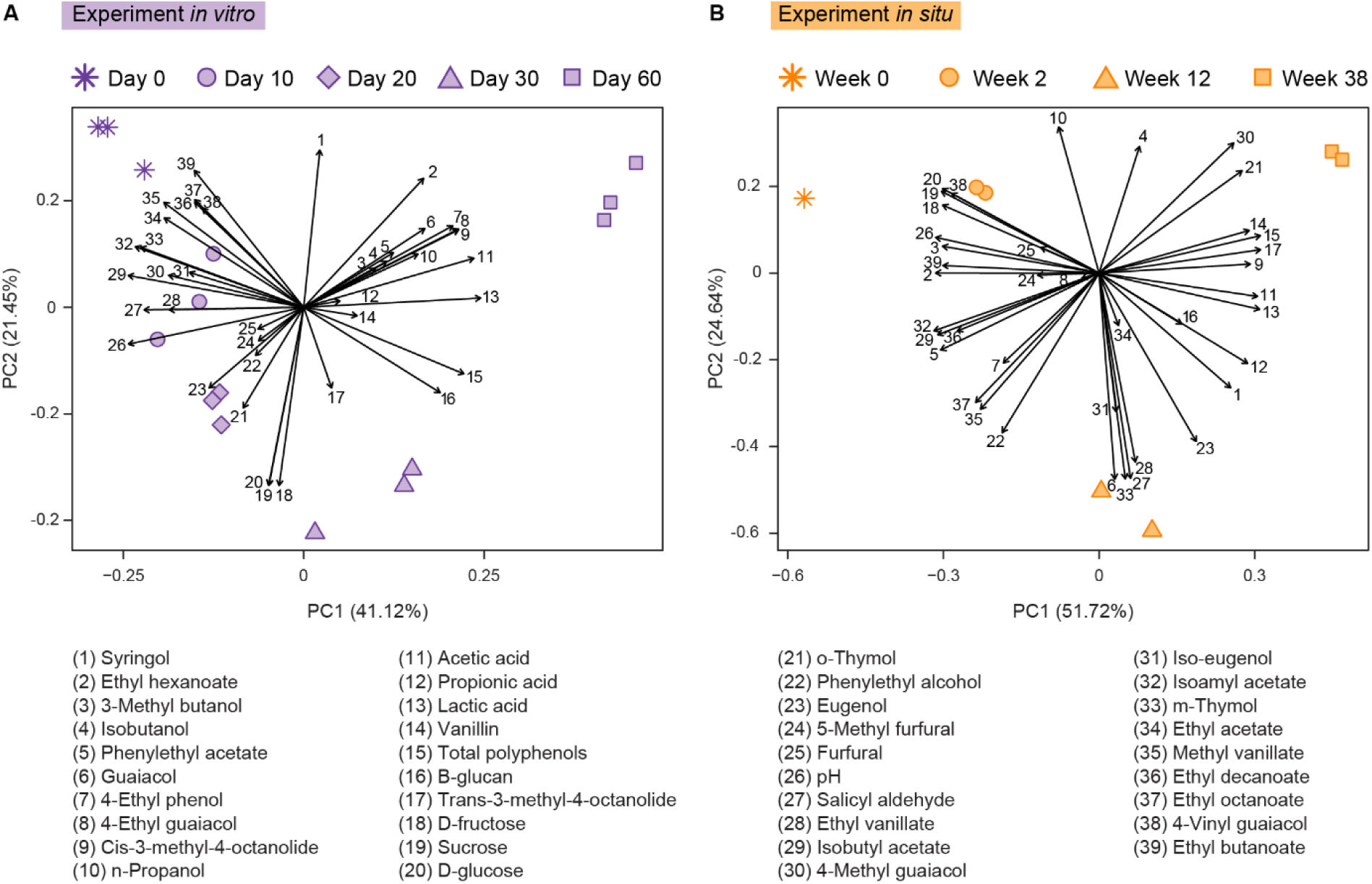
Principal component analysis (PCA) visualizing the differences in chemical composition of beer samples taken at different time points (indicated by different symbols) throughout (A) a 60-day maturation period of beer in an *in-vitro* system (*n* = 3) in which a synthetic microbial community of four bacterial (*Acetobacter malorum, Gluconobacter oxydans, Lactobacillus brevis*, and *Pediococcus damnosus*) and four fungal species (*Brettanomyces bruxellensis, Candida friedrichii, Pichia membranifaciens* and *Saccharomyces cerevisiae*) was pitched, and (B) an *in-situ* barrel-ageing experiment (*n* = 2) of 38 weeks. The same beer was used in both experiments. However, whereas the barrels were filled with non-filtered beer that still contained the *S. cerevisiae* strain from primary fermentation, the beer used in the *in-vitro* experiment was filter-sterilized. The chemical variables are presented as vectors. The smaller the distance between two data points, the more similar the chemical composition.

In general, similar changes in beer chemistry are observed in both experiments (Fig. 5; Fig. S3 and Tables S4 and S5, Supplementary Information). In both experiments pH decreased over time, while the concentrations of organic acids like acetic acid, lactic acid and propionic acid increased (Fig. 5). However, pH remained higher and the obtained lactic acid concentration remained lower in the *in-vitro* system than in the barrels, while a similar acetic acid concentration was reached in both experiments (Fig. 5). Further, sugar concentrations decreased throughout maturation, reaching slightly (although not significantly) lower concentrations in the *in-vitro* system than in the barrels at the end of the experiments. In contrast, despite that the β-glucan concentration increased in both experiments, concentrations measured in samples of the *in-vitro* system were more than 100 times higher than in the *in-situ* experiment (Fig. 5). A similar trend is observed for 4-ethyl guaiacol, 4-ethyl phenol and guaiacol with significantly higher concentrations obtained in the *in-vitro* system. Also, the concentration of cis-3-methyl-4-octanolide was higher after 60 days of maturation in the *in-vitro* system than after 38 weeks of ageing in barrels, although the difference in concentration was not significant. Additionally, concentrations of total polyphenols and vanillin increased throughout maturation, reaching significantly higher levels of total polyphenols and significantly lower levels of vanillin in the *in-vitro* system after 60 days of wood-ageing (Fig. 5). In contrast, furfural and 5-methyl furfural concentrations increased during ageing in the 225-liter barrels, whereas their concentrations first increased in the *in-vitro* system, but decreased after 10 days, reaching significantly lower concentrations of furfural at the end of the experiment in the *in-vitro* system than in the wooden barrels. Finally, the concentration of most esters decreased during ageing in both experiments, with comparable concentrations at the end of the experiments (Fig. 5). For the exact differences in beer chemistry between beer ageing in the *in-vitro* system and in barrels, the reader is referred to Tables S4 and S5 (Supplementary Information) and Fig. S3 (Supplementary Information).

**Figure 5:**
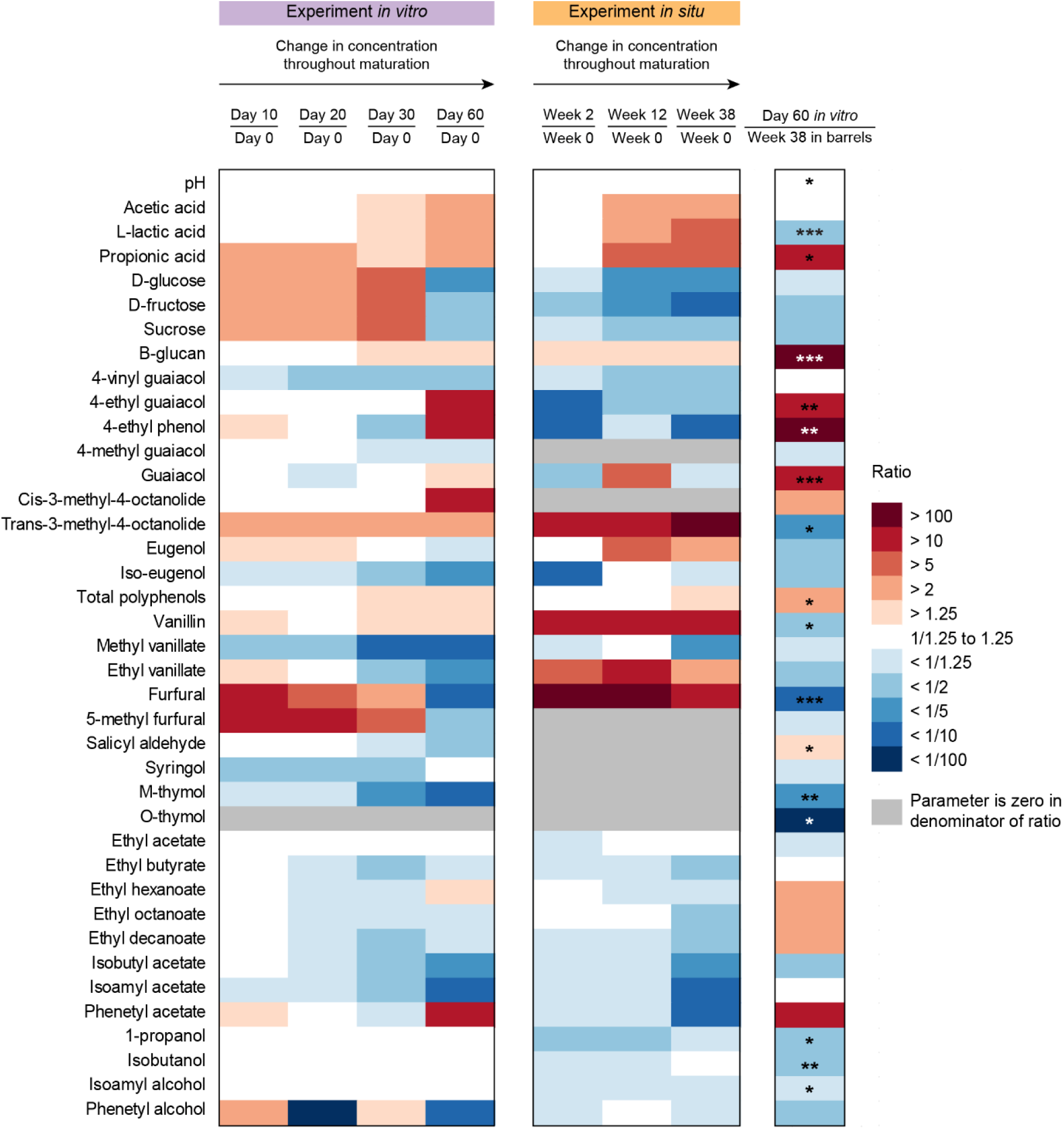
Heatmap visualizing the ratio between the concentration of the different chemical compounds at several time points throughout wood maturation, in comparison to the concentrations measured at the start, throughout a 38-week industrial-scale barrel-ageing experiment *in situ* (*n* = 2) and a 60-day *in vitro* wood maturation experiment (*n* = 3) in which a synthetic microbial community was pitched containing four bacteria (*Acetobacter malorum, Gluconobacter oxydans, Lactobacillus brevis* and *Pediococcus damnosus*) and four fungi (*Brettanomyces bruxellensis, Candida friedrichii, Pichia membranifaciens*, and *Saccharomyces cerevisiae*). Colours indicate whether the concentration of the chemical compounds increased (red) or decreased (blue) over time. Additionally, the ratio between the concentrations measured at the last time point of both experiments (right column) is shown, providing insight into the differences in concentration at the end of the maturation in both systems. In this case, also Welch t-tests were performed to test whether or not differences were significant. *P*-values obtained in the t-tests are presented in the cells: *\* p* < 0.05, ** *p* < 0.01, *** *p* < 0.001. Empty cells indicate that no significant differences (*p* ≥ 0.05) were obtained.

A main advantage of using the *in-vitro* system is that it easily allows to include more biological replicates as well as negative controls in which no microbes interfere with the process. Therefore, the *in-vitro* system enables an additional analysis looking at the effects of the pitched synthetic community in comparison to wood-ageing without any microorganisms (Fig. 6). As shown in Fig. 6, also wood-ageing without microorganisms substantially altered beer chemistry, including pH and concentrations of 4-vinyl guaiacol, 4-methyl guaiacol, methyl vanillate, ethyl vanillate, salicyl aldehyde, ethyl acetate, ethyl decanoate and isoamyl acetate which decreased significantly, while concentrations of acetic acid, D-glucose, D-fructose, sucrose, total polyphenols and vanillin increased significantly. When zooming in on the differences between the negative controls and samples that contained the synthetic microbial community (Fig. 6), after 60 days of maturation pH and concentrations of sugars, 4-vinyl guaiacol and vanillin were significantly lower in samples that contained microorganisms, whereas the concentrations of furfural and 5-methyl furfural were also more than 100 times lower in those samples, albeit not significantly. In contrast, concentrations of acetic acid, lactic acid, 4-ethyl guaiacol, 4-ethyl phenol, cis-3-methyl-4-octanolide, syringol, and ethyl hexanoate were significantly higher in beer pitched with microbes (Fig. 6).

**Figure 6:**
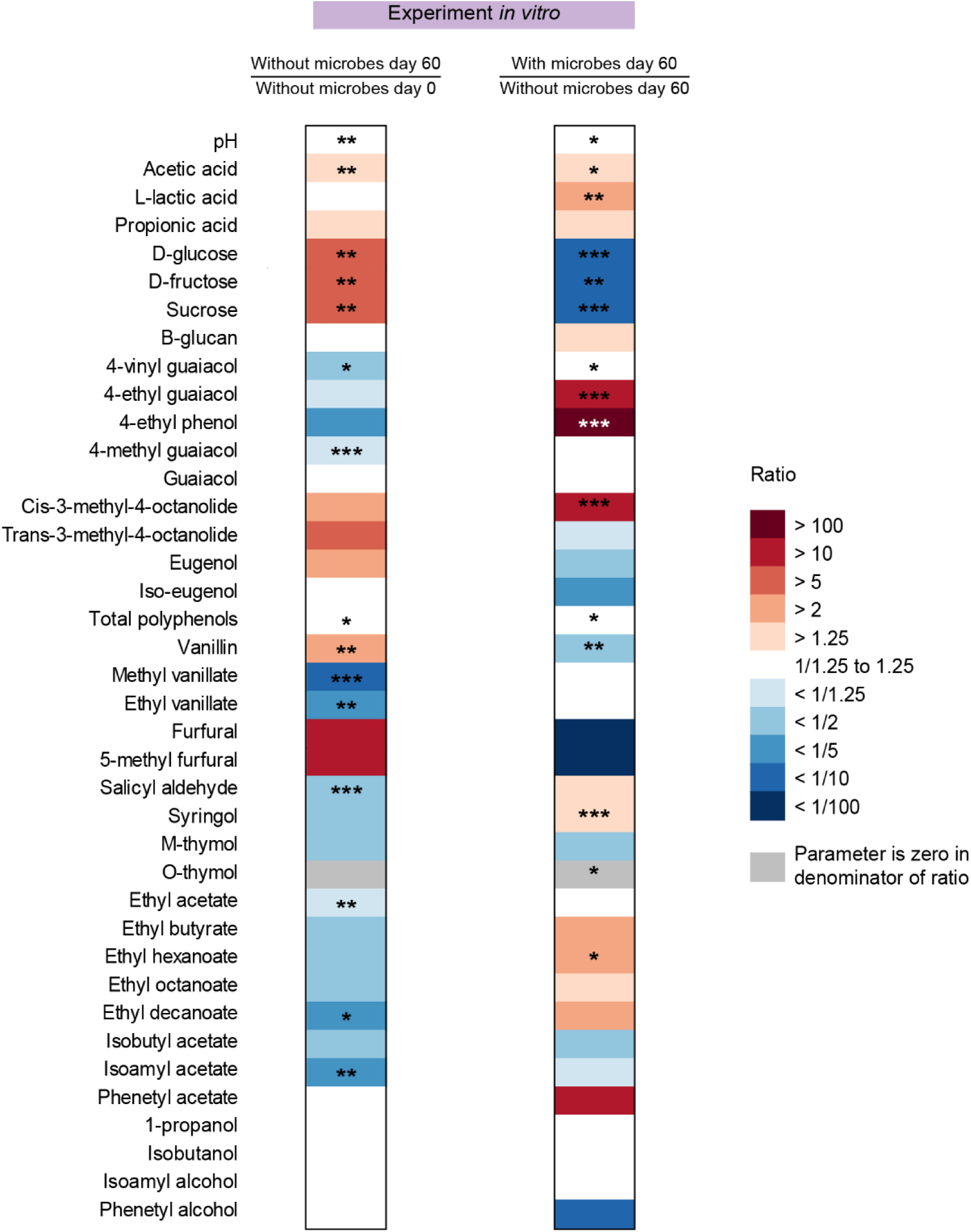
Heatmap based on beer chemistry assessed during a 60-day *in vitro* wood maturation experiment (*n* = 3) in which a synthetic microbial community was pitched containing four bacteria (*Acetobacter malorum, Gluconobacter oxydans, Lactobacillus brevis* and *Pediococcus damnosus*) and four fungi (*Brettanomyces bruxellensis, Candida friedrichii, Pichia membranifaciens* and *Saccharomyces cerevisiae*). The heatmap visualizes the ratio between the concentrations of different chemical compounds measured in the negative controls (without microbial community) at day 60 and the concentrations measured at day 0 (left column), as well as the ratio between concentrations measured in samples containing a microbial community at day 60 and concentrations in the negative controls at day 60 (right column). Whereas the left column indicates the impact of wood-ageing without microbes, the right column reveals the effect of the pitched microbial community. Welch t-tests were performed to test whether or not differences were significant. *P*-values obtained in the t-tests are presented in the cells: *\* p* < 0.05, ** *p* < 0.01, *** *p* < 0.001. Empty cells indicate that no significant differences (*p* ≥ 0.05) were obtained.

## 4. Discussion

In this study, we developed a lab-scale, experimentally tractable system to overcome the various challenges raised in studying industrial-scale barrel-ageing. Barrel-ageing was mimicked in a 0.5 liter weck jar with a wooden disk, while maintaining the ratio between the beer volume and wood surface area found *in situ*. Due to the small dimensions of the system, many jars can be simultaneously incubated in controlled experimental conditions, enabling an in depth study of the maturation process that goes well beyond any observational study performed previously. To speed up the process and increase reproducibility, the complexity of the natural microbial communities was reduced to a number of key microbes pitched into the simplified model system. The synthetic community was constructed based on a number of criteria. Firstly, all species involved should be culturable and not pose any health risks. Furthermore, the species selected should represent major groups associated with the process of barrel-ageing, including acetic acid bacteria, lactic acid bacteria, and conventional and ‘wild’ (souring) fungi that dominate the community (Bossaert *et al*., 2021a; 2021b). In addition, the synthetic community should possess a few microbes representing the background flora present, which may become important when less-stringent conditions are applied (Bossaert *et al*., 2021a). To meet these criteria, a synthetic community was composed containing four bacterial and four fungal strains isolated from barrel-aged beer. More specifically, the synthetic community contained the acetic acid bacteria *A. malorum* and *G. oxydans*, the lactic acid bacteria *L. brevis* and *P. damnosus, S. cerevisiae* and the wild yeasts *B. bruxellensis, C. friedrichii* and *P. membranifaciens*. In our previous experiments *G. oxydans* and *C. friedrichii* were particularly found at the beginning of the process, while the other species were all found to dominate the community when maturation time progressed (Bossaert *et al*., 2021a; 2021b). Moreover, importantly, all of them have been shown to contribute to the aromas produced during barrel-ageing of beer. Acetic acid bacteria like *A. malorum* and *G. oxydans* generally produce acetic acid, gluconic acid, glucuronic acid, and ethyl acetate, and can sometimes produce acetoin from lactic acid (De Roos *et al*., 2018; Kashima *et al*., 1998; Moens *et al*., 2014). Lactic acid bacteria like *L. brevis* and *P. damnosus* mainly contribute lactic acid, although certain strains can also produce 4-ethyl guaiacol and 4-ethyl phenol, or exhibit α-glucosidase activity (Couto *et al*., 2006; De Cort *et al*., 1994; Giraffa *et al*., 2010). Additionally, studies about the malolactic fermentation of wine have revealed an interesting, yet intricate interaction between lactic acid bacteria and wood-derived compounds such as vanillin and oak lactones (Bastard *et al*., 2016; Bloem *et al*., 2007; de Revel *et al*., 2005). In fact, when malolactic fermentation of wine was performed by *Œnococcus œni* in contact with wood, higher concentrations of oak lactones, eugenol, iso-eugenol and vanillin were measured (de Revel *et al*., 2005), whereas the concentration of furfural was found to be lower (Bastard *et al*., 2016). Moreover, in culture medium spiked with vanillin, higher cell densities of *Œ. œni* were observed in comparison to the medium without vanillin (de Revel *et al*., 2005). Further, the conventional yeast species *S. cerevisiae* is known to be an efficient fermenter and produces fruity and floral aromas that bode well with the sour flavour profile of the beer (De Roos and De Vuyst, 2019). *B. bruxellensis* produces ethanol and acetic acid, as well as 4-ethyl guaiacol, 4-ethyl phenol, and various ethyl esters including ethyl acetate, i.e. compounds that are known to contribute to the ‘funky’ character of traditional Belgian Lambic beers (De Roos and De Vuyst, 2019). Furthermore, certain *B. bruxellensis* strains exhibit α- and β-glucosidase activity capable of hydrolysing glycosides, which could potentially contribute to the beer aroma or allow the yeasts to thrive and survive in the barrels (Crauwels *et al*., 2015; Steensels *et al*., 2015). *Pichia* spp. can also produce ethanol, lactic acid and ethyl acetate among other acetate esters like isoamyl acetate, and can create slightly sour beers with fruity, aromatic flavours (Holt *et al*., 2018; Osburn *et al*., 2018). Additionally, certain strains of *P. membranifaciens* show activity of β-glucosidase and 1,4-β-xylosidase that can be advantageous to survive in wooden barrels as these enzymes play a role in the breakdown of cellobiose (cellulose) and xylans (hemicellulose), respectively (López *et al*., 2015). Finally, *Candida* spp. can produce ethanol, acetic acid, glycerol and in a few strains β-glucosidase activity has been observed as well (Englezos *et al*., 2015; Estela-Escalante *et al*., 2016). As a result, all species selected for the synthetic microbial community play a specific role within the community and throughout wood maturation, and contribute to the complex, layered flavour profile of the resulting sour beers.

To monitor the different strains, species-specific qPCR assays were developed. A major advantage of qPCR over an amplicon sequencing approach is that it provides fast and high-throughput detection and absolute quantification of target DNA sequences, while the amplicon sequencing approach results in relative abundance profiles. As such, the qPCR assays are particularly useful to monitor the different strains over time. The assays were highly reliable and did not yield a single cross-reaction with any of the strains tested. Furthermore, the assays allowed gene copy quantifications over a range of at least nine orders of magnitude, with LOQ values ranging between 0.81 and 1.88 log gene copies per µl DNA extract. These results are in agreement with previous studies, illustrating the high sensitivity of our qPCR assays (Bokulich *et al*., 2012; Bonk *et al*., 2018; Justé et al., 2008). Moreover, measured gene copy numbers of the microbial populations in the *in-vitro* experiment were confirmed by total bacterial and fungal qPCR assays as well as by plating on different cultivation media (Fig. S4, Supplementary Information), illustrating the usefulness of the qPCR assays.

Microbial and chemical analysis of all samples taken in the *in-vitro* experiment illustrate the great reproducibility of the system. Indeed, biological replicates yielded highly similar microbial community (Fig. 2) and chemical (Fig. 4) profiles. This reproducibility is one of the major advantages of an *in-vitro* system with a synthetic microbial community, as it is a crucial element in studying the impact of individual abiotic factors on the community dynamics and associated chemical changes, without the interfering variability that is often associated with natural communities. Moreover, studies that aim to unravel the mechanisms of community assembly in a particular study system, like cheese rinds (Wolfe *et al*., 2014), broad bean paste fermentations (Jia *et al*., 2020) or Chinese light-aroma-type liquor fermentations (Wang *et al*., 2019), rely on this reproducibility to facilitate a mechanistic understanding of the structure and function of the microbiota and to later use this knowledge to improve food fermentation processes on an industrial scale to guarantee a consistent, desirable end quality. When comparing the microbial populations in the *in-vitro* system and the industrial-scale wooden barrels, no significant differences were detected between the bacterial and fungal populations at the first and last sampling point during wood-ageing *in vitro* (day 60) and on an industrial scale (week 38). This suggests that similar population dynamics were found in the experiments on both scales. However, as only a limited number of barrels (i.e. biological replicates) could be included in the industrial-scale barrel-ageing, the number of permutations that could be performed in the perMANOVA analysis was restricted, which may have affected the significance of the results. Indeed, especially for fungi, the NMDS plot seems to suggest differences in populations as data points for the *in-vitro* experiment and the *in situ* experiment were plotted distantly from each other. When zooming in on the different microorganisms, in both experiments 16S rRNA gene copy numbers of *L. brevis* increased reaching approximately 7 log gene copy numbers at the end of the experiments, whereas *G. oxydans* gene copy numbers remained low over the entire experimental period. In contrast, gene copy numbers of *A. malorum* and *P. damnosus* remained low in the *in-vitro* system, whereas they reached relatively high gene copy numbers in the barrels. Although the reason for this discrepancy is still unknown, it is reasonable to assume that strain differences may have played a role. In other words, strains selected for the synthetic community were isolated from ‘beer 2’ in Bossaert *et al*. (2021a) and not from the beer under investigation in this study, even though *A. malorum* and *P. damnosus* also occurred there. Therefore, the strains included in the experiment *in vitro* could exhibit different phylogenetic characteristics than the strains detected *in situ*, and they may not be adequately adapted to growth conditions posed by the beer in this study (Matsutani *et al*., 2012; 2020; Shani *et al*., 2021; Snauwaert *et al*., 2015). Further, ITS copy numbers of *B. bruxellensis, P. membranifaciens* and *S. cerevisiae* increased throughout maturation in both experiments, albeit not to the same extent. Whereas 7.74 ± 0.02, 6.85 ± 0.17 and 5.32 ± 0.08 log ITS copy numbers were reached *in vitro* for *B. bruxellensis, P. membranifaciens* and *S. cerevisiae*, respectively, only *S. cerevisiae* reached a similar level of 5.80 ± 0.19 log ITS copy numbers *in situ*, while *B. bruxellensis* and *P. membranifaciens* grew to 1.0 ± 0.37 and 2.7 ± 1.41 log ITS copies in the barrels. As unfiltered beer was used in the barrels (Bossaert *et al*., 2021a), the *S. cerevisiae* strain used in primary fermentation was still present in 5.53 log ITS copies at the start of the maturation *in situ*, whereas the beer used *in vitro* was filter-sterilized before use and all microbes were pitched in equal cell densities. Therefore, *S. cerevisiae* was not given a head start over the other yeast strains in the *in-vitro* system, which may be the reason why *B. bruxellensis* and *P. membranifaciens* could grow to higher densities *in vitro*, while their growth was restricted in the barrels. Accordingly, in the wooden barrels, the *S. cerevisiae* strain may have had the chance to consume any residual nutrients before the other organisms were given the chance. Strikingly, copy numbers of *C. friedrichii* remained under the detection limit in both experiments, suggesting that this yeast is not able to thrive under the conditions applied. In contrast, growth of this *C. friedrichii* strain was observed in several of the conditions applied in the pilot screening (data not shown). Likewise, in the synthetic community developed by Wang *et al*. (2019), the *Candida* strain was able to grow to a high relative abundance in the presence of *Lactobacillus, Saccharomyces* and *Pichia*. Further, the impeded growth of *A. malorum* and/or *P. damnosus* in the *in-vitro* system could be due to one of the yeasts that is abundantly present in the *in-vitro* system but not in barrels, i.e. *B. bruxellensis* or *P. membranifaciens*. Indeed, these yeasts could have altered the beer medium to restrict growth of these bacterial strains, e.g. by lowering pH, by producing organic acids or by the consumption of nutrients that are needed for growth of *A. malorum* or *P. damnosus* (Hibbing *et al*., 2010; Liu *et al*., 1996; Nguyen *et al*., 2015; Niu *et al*., 2016).

Also beer chemistry of both experiments was not significantly different at the first and last sampling point during maturation. However, due to the limited number of biological replicates included in the industrial-scale experiment, the statistical power of the perMANOVA may not have been sufficient to establish significant differences. In contrast, maturation time significantly affected beer chemistry during ageing in the *in-vitro* system as well as in the *in-situ* experiment. Indeed, chemical parameters varied throughout maturation, but followed similar trends in both experiments. More specifically, in both experiments, pH and concentrations of sugars, 4-vinyl guaiacol and most esters were lower at the end of the maturation period, while the concentration of organic acids increased throughout wood-ageing. Nevertheless, the final concentration reached after 60 days of ageing in the *in-vitro* system was for several compounds significantly different from the concentration obtained after 38 weeks of barrel-ageing. In fact, the pH was higher and the concentration of lactic acid was lower in beer aged in the *in-vitro* system. Further, the concentrations of β-glucan, 4-ethyl guaiacol, 4-ethyl phenol and total polyphenols were significantly higher, and concentrations of vanillin, furfural and M- and O-thymol were significantly lower in the experiment *in vitro* than in the experiment *in situ*. In fact, many of these compounds were significantly linked to the microbial community *in vitro*, including pH, lactic acid, 4-ethyl guaiacol, 4-ethyl phenol, total polyphenols, vanillin and O-thymol among others, whereas also the other compounds like β-glucan, furfural and M-thymol seemed affected by the microbes, albeit not significantly (Fig. 5 and 6). Therefore, it is plausible that differences in beer chemistry between both experiments were caused by differences in the microbial community.

One of the main advantages of the *in-vitro* system is that it is small, robust, reproducible and easy-to-manipulate. Therefore, it can be easily altered to evaluate the impact of diverse parameters of interest, including the screening of different wood species, different beer styles, pairing beer styles with wood types, as well as the screening of promising microbial strains for sour beer production via barrel-ageing. Further, the system pitched with the synthetic microbial community is perfectly equipped to study the mechanisms that underlie microbial successions and chemical changes during barrel-ageing of beer, which could help develop process-control strategies and improve the predictability and consistency of barrel-aged sour beers. Additionally, using the *in-vitro* system developed, it becomes much more feasible to include more replicates and the necessary controls to reinforce statistical conclusions and study the individual effects of the microbial community, the beer, the wood, and study different parameters simultaneously. Moreover, this allows more in-depth research on how external, abiotic factors like beer properties affect the temporal dynamics in microbial community composition and beer chemistry in a statistically meaningful manner.

## 5. Conclusion

The lab-scale, experimentally tractable system developed in this study was proven to be a robust and reproducible system that could overcome the challenges raised by studying barrel-ageing of beer on an industrial scale. The *in-vitro* system was equipped with a synthetic community composed of four bacterial species (*A. malorum, G. oxydans, L. brevis* and *P. damnosus*) and four fungal species (*B. bruxellensis, C. friedrichii, P. membranifaciens* and *S. cerevisiae*) that represented key microbes previously identified in experiments with 225-liter barrels. Each microorganism can be monitored by a species-specific qPCR assay, allowing accurate strain detection and quantification. While not completely perfect, the temporal dynamics in microbial community composition and beer chemistry observed after barrel-ageing for 38 weeks in 225-liter barrels could be mimicked throughout 60 days of ageing *in vitro*. For this reason, the *in-vitro* system opens the door to performing more in-depth research about the intricate interactions between microbes, wood and maturing beer, as well as the mechanisms of microbial community assembly and the impact of external, abiotic factors like beer properties on the temporal dynamics in the microbial community and changes in beer chemistry.

## Supporting information

Standard curves qPCR genomic DNA

Standard curves qPCR amplicon DNA

Line plots chemistry

Line plots chemistry2

Plate counts

## 7. Data availability

The data used in this study (qPCR and chemical data) are available in Supplementary Information. Sequences of the 16S rRNA gene (bacteria), and the 28S rRNA gene or the ITS1-5.8S rRNA-ITS2 region (fungi) of the microbial strains used in this study have been deposited in GenBank under the following accession numbers: OM432146 (*A. malorum*); OM432147 (*G. oxydans*); OM432145 (*L. brevis*); OM432144 (*P. damnosus*); OM432156 (*S. cerevisiae*); OM432150 (*B. bruxellensis*); OM432151 (*P. membranifaciens*); and OM432152 (*C. friedrichii*).

## 8. Supplementary data

**Table S1:** Data log of temperature, humidity and dew point over the course of the experiment.

**Table S2:** Specifications of the chemical analysis protocols.

**Table S3:** qPCR data expressed as gene copy numbers per μl DNA.

**Table S4:** pH and concentration of carbohydrates, ethanol and fermentation products.

**Table S5:** Concentrations of wood and hop compounds.

**Figure S1**: qPCR standard curves created using 10-fold dilutions of genomic DNA.

**Figure S2**: qPCR standard curves created using 10-fold dilutions of DNA amplicons.

**Figure S3**: Temporal changes in beer chemistry during wood-ageing of beer in two experimental set-ups: (i) 60 days of wood maturation in an *in-vitro* system pitched with four bacteria (*Acetobacter malorum, Gluconobacter oxydans, Lactobacillus brevis* and *Pediococcus damnosus*) and four fungi (*Brettanomyces bruxellensis, Candida friedrichii, Pichia membranifaciens* and *Saccharomyces cerevisiae*), and (ii) 38 weeks of barrel-ageing *in situ*. Data are presented as the average of biological controls (*n* = 3 *in vitro*, and *n* = 2 *in situ*) and the error bars represent the associated standard error of the mean. Displayed parameters: (A) pH, (B) acetic acid, (C) lactic acid, (D) D-glucose, (E) D-fructose, (F) sucrose, (G) β-glucan, (H) 4-vinyl guaiacol, (I) 4-ethyl guaiacol, (J) 4-ethyl phenol, (K) eugenol, (L) iso-eugenol, (M) cis-3-methyl-4-octanolide, (N) trans-3-methyl-4-octanolide, (O) total polyphenols, (P) vanillin, (Q) furfural, (R) 5-methyl furfural, (S) ethyl acetate, (T) isoamyl acetate. For a detailed overview of the different chemical parameters measured in this study, the reader is referred to Tables S4 and S5 (Supplementary Information).

**Figure S4**: Temporal dynamics in microbial community composition assessed via cultivation on different media at several time points during *in vitro* wood maturation. The *in-vitro* system was pitched with four bacteria (*Acetobacter malorum, Gluconobacter oxydans, Lactobacillus brevis* and *Pediococcus damnosus*) and four fungi (*Brettanomyces bruxellensis, Candida friedrichii, Pichia membranifaciens* and *Saccharomyces cerevisiae*).

