## Supplementary figures and images for "Development of a tractable model system to mimic wood-ageing of beer on a lab scale"

### Line plots chemistry

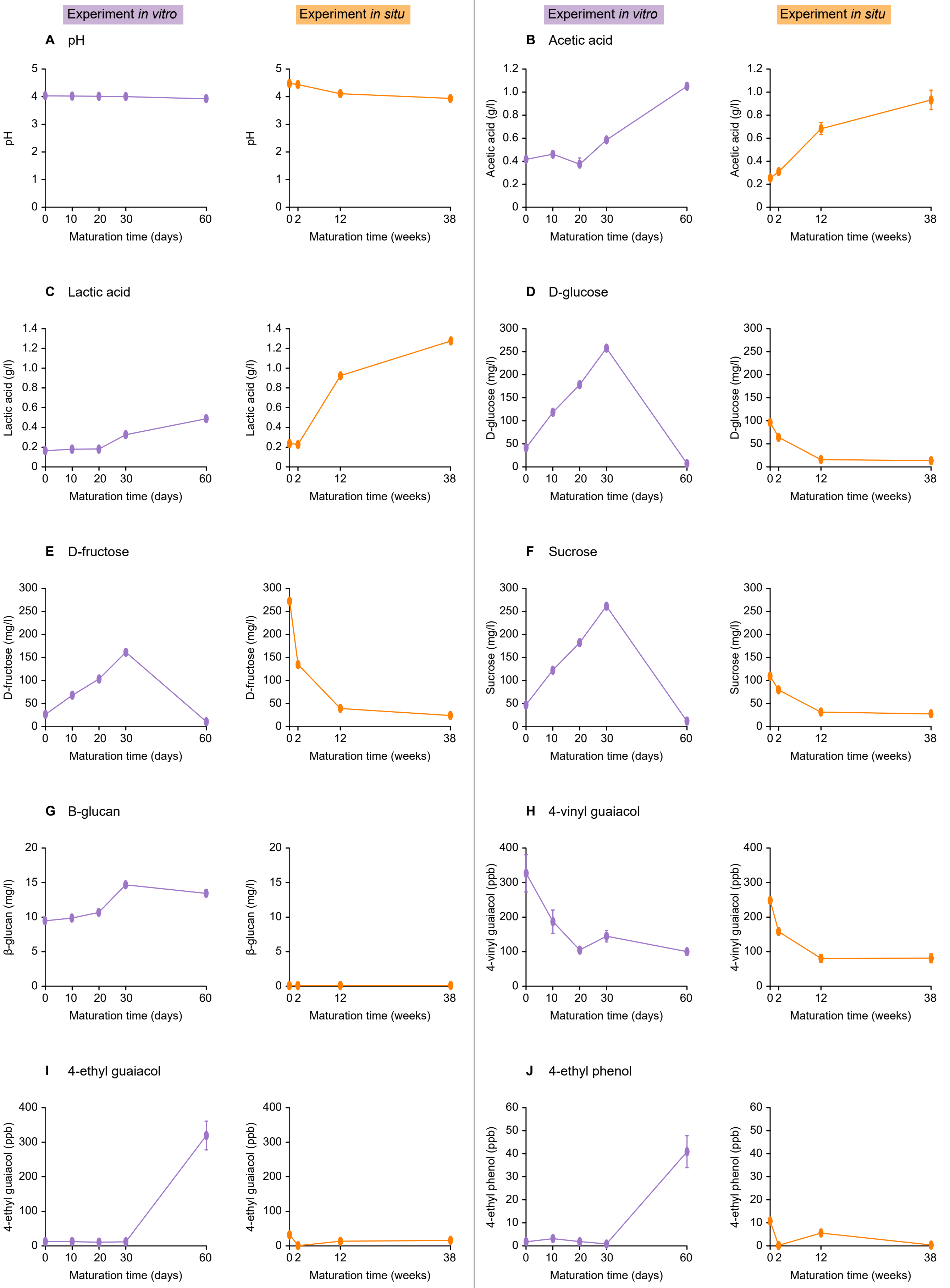

### Line plots chemistry2

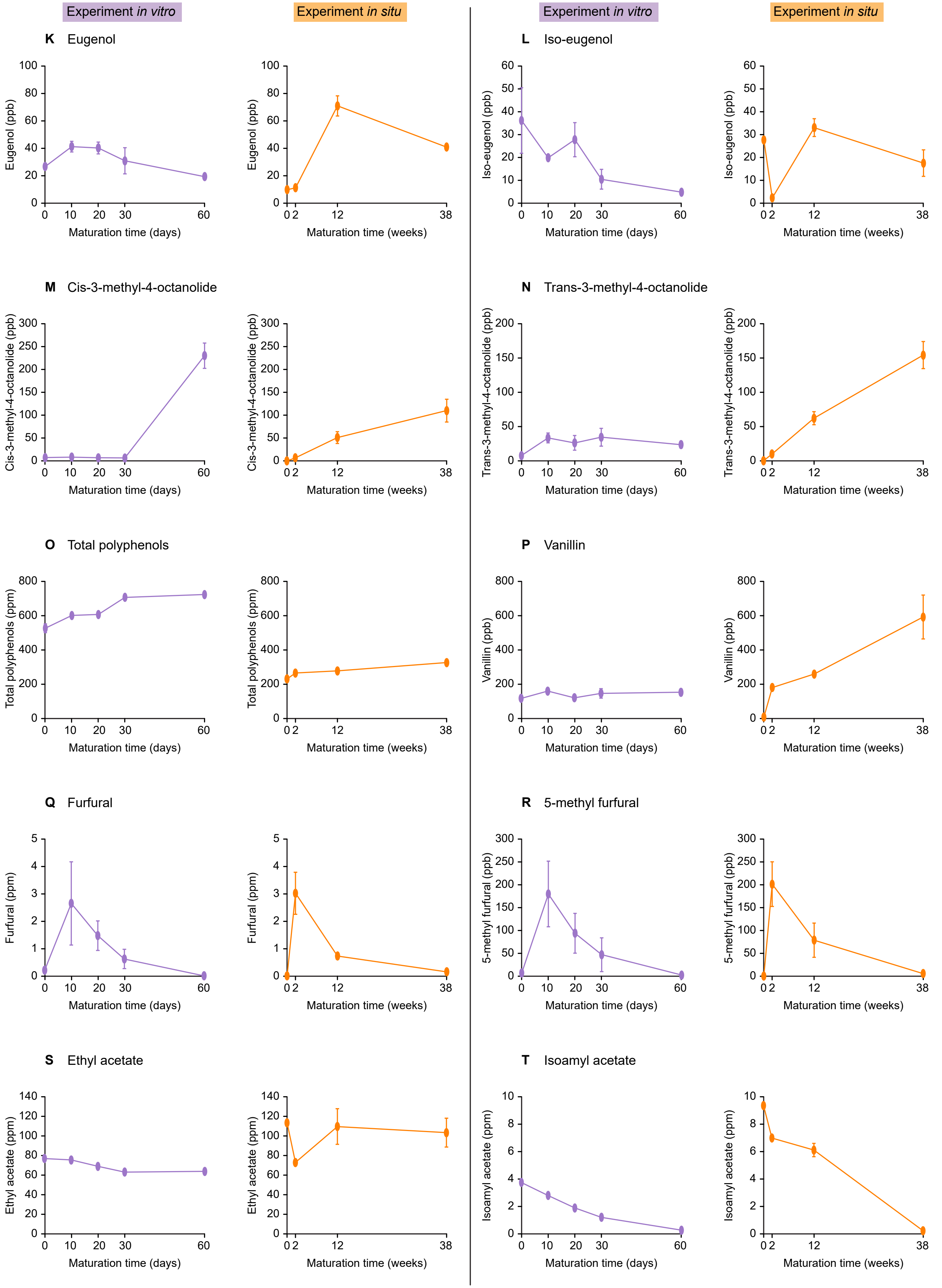

### Plate counts

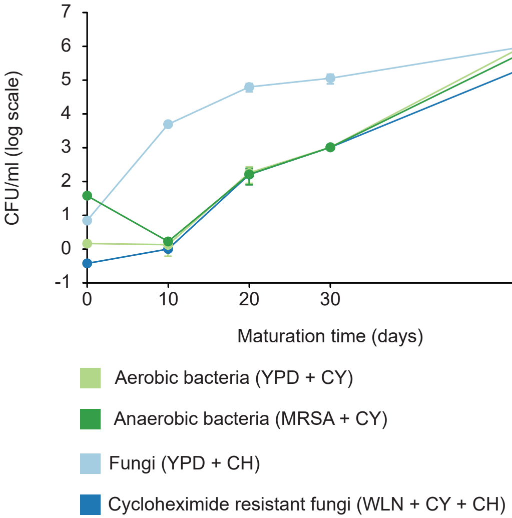

### Standard curves qPCR amplicon DNA

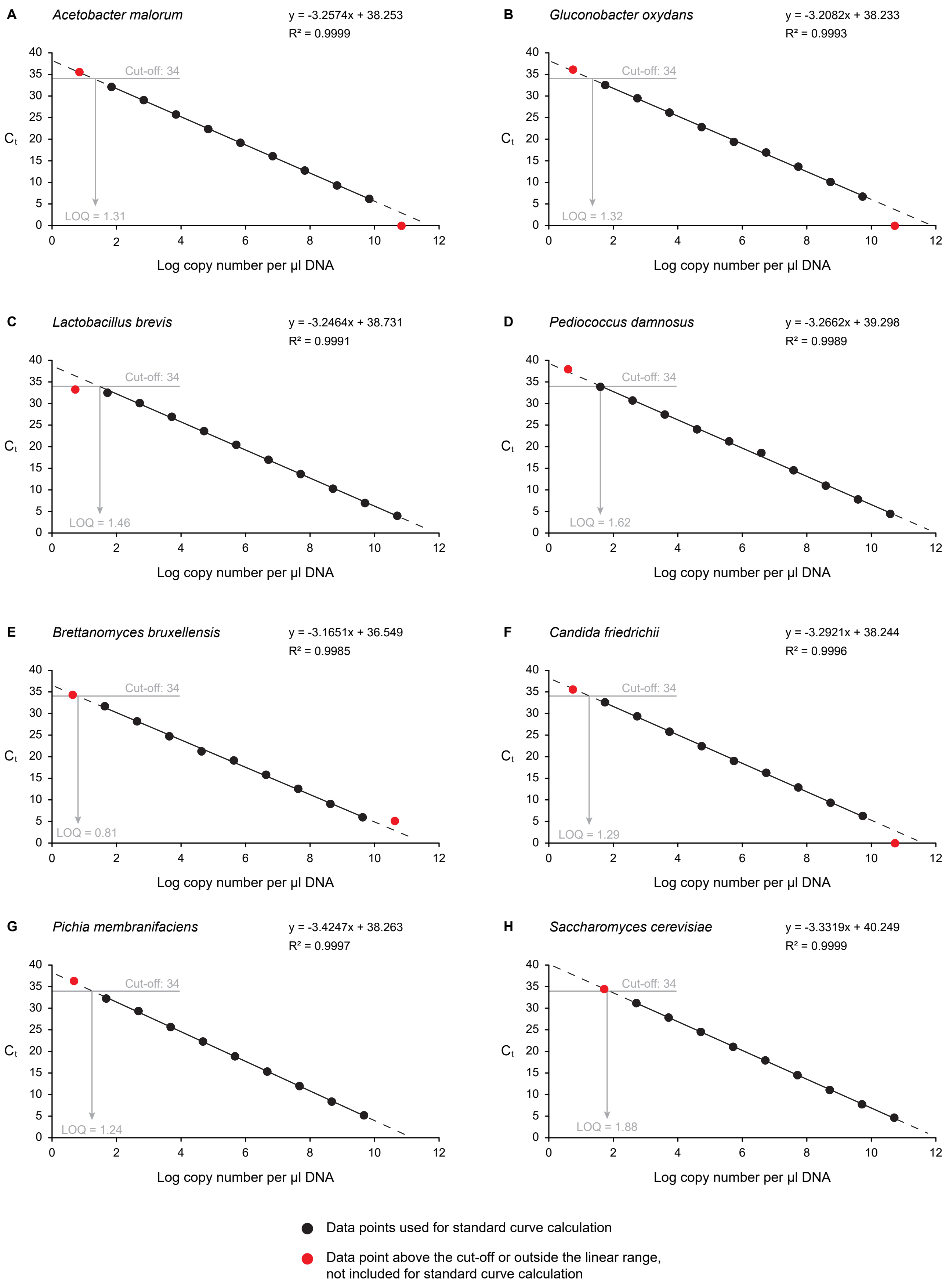

### Standard curves qPCR genomic DNA

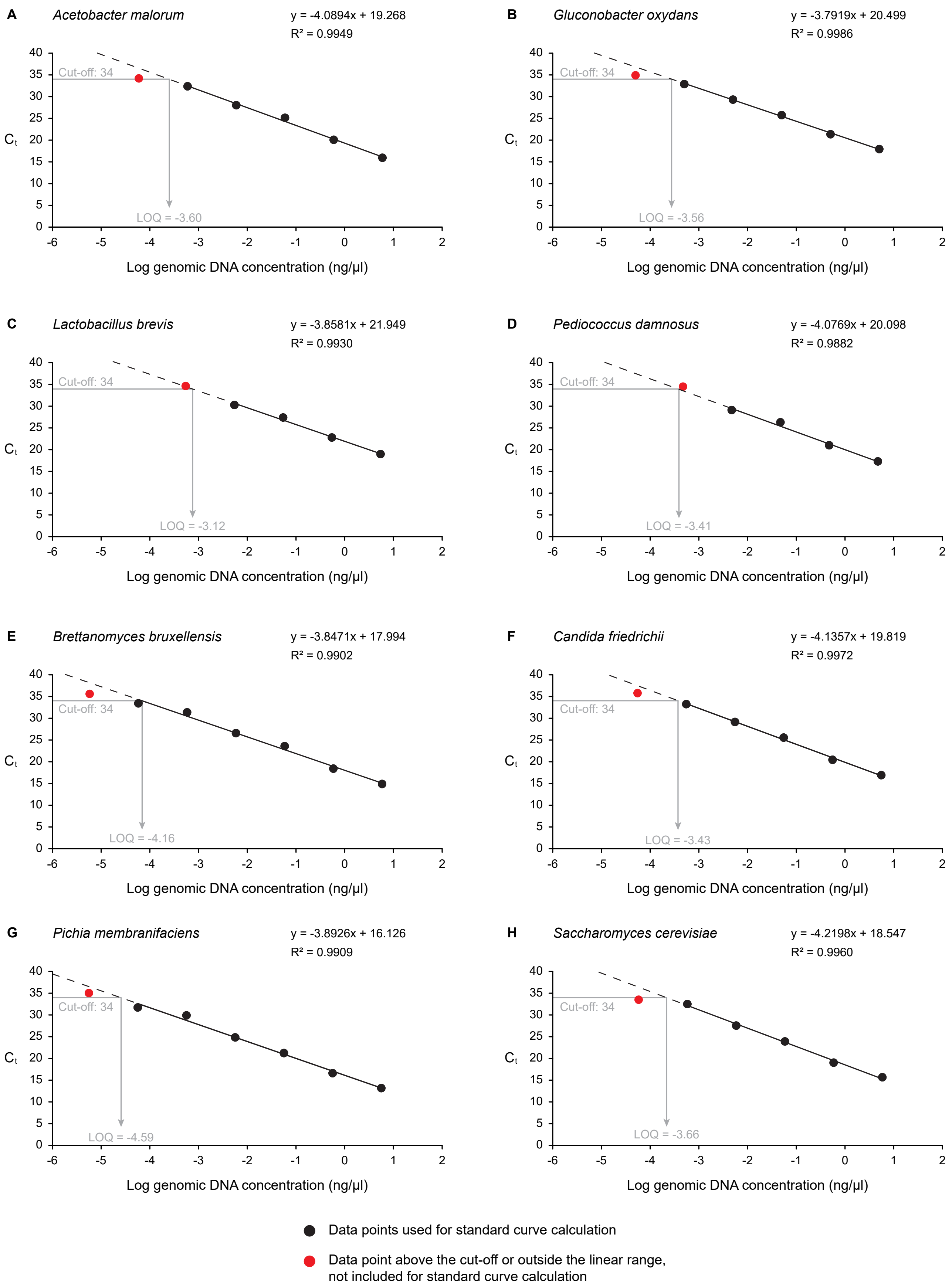
